# The two-step purification method ViREn identifies a single NSUN6-mediated 5-methylcytosine modification promoting dengue virus RNA genome turnover

**DOI:** 10.1101/2025.03.17.643699

**Authors:** Chia Ching Wu, Isabel S Naarmann-de Vries, Jonas Hartmann, Zarina Nidoieva, Kevin Kopietz, Virginie Marchand, Zeynep Özrendeci, Doris Lindner, Sophia Schelchshorn, Sophia Flad, Michaela Frye, Nina Papavasiliou, Tanja Schirmeister, Georg Stoecklin, Johanna Schott, Yuri Motorin, Francesca Tuorto, Christoph Dieterich, Mark Helm, Alessia Ruggieri

## Abstract

Chemical modifications on cellular and viral RNAs are new layers of post-transcriptional regulation of cellular processes including RNA stability and translation. Although advances in analytical methods have improved detection sensitivity, the precise mapping of RNA modifications at single-base resolution remains challenging. Especially for low abundant viral RNAs extracted from infected cells, requirements for sensitivity and purity limit accuracy and reproducibility. Here we report the two-step method ViREn for the enrichment of the genomic RNA (gRNA) of dengue virus (DENV), a positive-sense single-stranded RNA virus. This approach enabled the preparation of gRNA with significantly increased purity and led to the identification of a high-confidence 5-methylcytosine (m^5^C) site in DENV gRNA, orthogonally validated by Illumina-based bisulfite sequencing and direct RNA sequencing by Nanopore Oxford Technologies. Strikingly, this m^5^C modification was exclusively detected in gRNA extracted from infected cells but not in gRNA extracted from viral particles. We identified NSUN6 as the host methyltransferase catalyzing this modification and demonstrated a role for m^5^C in regulating DENV gRNA turnover. ViREn thus enables the mapping of m^5^C on low abundance viral gRNA with unprecedented precision and sensitivity and facilitates mechanistic studies into the role of RNA modification in virus replication.

## INTRODUCTION

Several classes of coding and noncoding RNA molecules carry chemical modifications (1–3), including messenger RNAs (mRNAs) at low abundance (4). Modifications can affect RNA structure and regulate cellular processes including splicing, nuclear export, stability, and translation (5). Recent advances of analytical methods have improved sensitivity for the detection of RNA modifications, e.g. using liquid chromatography–tandem mass spectrometry (LC-MS/MS) (6). Similarly, the combined use of antibody-enriched or chemical-treated RNA with RNA sequencing (RNA-Seq) approaches has improved the resolution and mapping of most abundant RNA modifications, such as N6-methyladenosine (m^6^A) and 5-methycytosine (m^5^C), at single-base resolution (7). However, these approaches still suffer from specific disadvantages, e.g. non-specific binding of antibodies, incomplete chemical conversion in the case of chemical treatments, low sensitivity for rare modifications, high material requirements, which affect accuracy and reproducibility. The development of direct RNA sequencing (DRS) by Nanopore Oxford Technologies (ONT) has paved the way for detecting RNA modifications in native single RNA molecules, particularly mRNAs, significantly contributing to our understanding of the dynamic epitranscriptome (8,9).

These advances in RNA-Seq based methods have also reinvigorated the search for RNA modifications in viral RNAs, particularly in the genome of positive-sense (+) single-stranded (ss) RNA viruses since modifications may directly influence their replication. The combined use of antibody-based purification and RNA-Seq has allowed the identification of multiple m^6^A-modified regions within the genome of several *Flaviviridae*, including dengue virus (DENV), Zika virus, Yellow Fever virus, West Nile virus and hepatitis C virus (HCV), suggesting a potentially conserved function across this virus family (10,11). In comparison, less is known about the m^5^C in the genome of *Flaviviridae*, probably due to its relatively low abundance. Furthermore, over 30 additional types of RNA modifications were identified in the genomic RNAs (gRNAs) of several RNA viruses by RNA affinity capture coupled to LC-MS/MS (12). However, some of the results have proven poorly reproducible, highlighting methodological shortcomings due mainly to low viral RNA abundance and contaminations by residual cellular RNAs such as highly modified transfer RNAs (tRNAs) and ribosomal RNAs (rRNAs). This has led to controversy over the positional mapping of modifications and underscored the necessity for orthogonal and functional validation (13,14).

DENV is a mosquito-borne human pathogen and a major public health burden. Dengue cases have seen an unprecedented increase in 2024, reaching 14 million infections and more than 10,000 dengue-related deaths worldwide^1^. The DENV gRNA is approximately 10.7 kilobases long, flanked by highly structured untranslated regions (UTRs) at the 5’ and 3’ ends. The single open reading frame of DENV gRNA encodes a polyprotein that is both co- and post-translationally cleaved to yield three structural and seven non-structural (NS) proteins involved in gRNA replication and viral particle production. The gRNA is capped at the 5’ end by the viral RNA-dependent RNA polymerase NS5 but lacks a polyadenylated tail at its 3’ end (15). The DENV replication cycle is exclusively cytoplasmic. During infection, the 5’ portion of the gRNA is degraded by the cellular 5’-3’ exonuclease XRN1, an essential component of the decapping-dependent RNA decay pathway (16). XRN1 stalls at compact RNA folds in the DENV gRNA 3’ UTR, generating small noncoding RNAs called subgenomic flaviviral RNA (sfRNA) (17,18). The cytosolic accumulation of sfRNA during infection is part of the viral strategy to counteract cellular antiviral responses in mosquito and vertebrate hosts (19–23), and contributes to viral pathogenicity (22–24).

Accurate identification and mapping of modified ribonucleotides have improved with advanced sequencing methods and basecalling algorithms. Nevertheless, the precise detection of RNA modifications remains affected by sequencing depth, the design of appropriate controls for algorithm training, and the number of biological replicates analyzed (25). Low purity and yield of viral RNA isolation protocols are thus the main obstacles to accurate modification detection. To overcome this limitation, we here developed a viral RNA enrichment method – called ViREn – for the in-depth sequencing and mapping of DENV gRNA modifications. ViREn is a two-step approach that separates DENV RNAs from most other cellular RNAs based on their sedimentation coefficient in sucrose gradients, followed by sequence-specific affinity capture. Using Illumina-based bisulfite sequencing (BSSeq) and DRS, we identified a single m^5^C site at position 1218 of the DENV gRNA with high confidence. This modification, exclusive to gRNA extracted from infected cells and undetectable in gRNA extracted from viral particles, is catalyzed by the cellular methyltransferase Nol1/Nop2/SUN 6 (NSUN6) and promotes DENV gRNA turnover.

## MATERIAL AND METHODS

### Cells

Huh7 cells, Huh7 NT cells and Huh7 NSUN6 KO cells were maintained at 37 °C, 5% CO^2^ in Dulbecco’s Modified Eagle Medium (DMEM, Gibco) supplemented with 10 % fetal calf serum (Capricorn), 1% nonessential amino acids, 100 U/ml penicillin and 100 μg/ml streptomycin (all from Gibco).

### Plasmids

The following plasmids were used and described elsewhere: pDVWSK601, harboring the full-length cDNA of DENV type 2 New Guinea C (26) (kindly provided by Andrew Davidson, University of Bristol, England) was used for production of DENV2 viral stocks and *in vitro* synthesis of unmethylated control RNAs for BSSeq and nanopore sequencing. pDVWSK601-LucUbi (27) harboring DENV NGC Firefly luciferase reporter virus was used for the characterization of DENV replication in Huh7 NSUN6 KO cells.

The plasmid pDVWSK601 C1218T was generated using pDVWSK601 as a template. The point mutation was introduced by site directed mutagenesis using overlap extension PCR. Two sets of degenerate PCR primers were: DENV-E_BsrGI_For: 5’-AGAGAAACCGCGTGTCGACTGTACAACAGCTGACAAAG-3’ and DENV-E_C1218T_Rev: 5’-CATCCTCTGTCCACCATGGAATCTTTGCAGACGAACCT-3’; DENV-E_C1218T_For: 5’-AGGTTCGTCTGCAAACATTCCATGGTGGACAGAGGATG-3’ and DENV-E_NheI_Rev: 5’-AGCTTTCTGGATAGCTGAAGCTTTGAAGGGGATTCTGG-3’. Amplicons were amplified using Phusion DNA polymerase (New England Biolabs, NEB), separated by agarose gel electrophoresis, purified from the gel using NucleoSpin Gel and PCR Clean-up (Macherey-Nagel). Equal amounts of each amplicon were used as template and combined by overlap extension PCR with primers DENV-E_BsrGI_For and DENV-E_NheI_Rev. PCR products were separated by agarose gel electrophoresis, purified, and subsequently digested with BsrGI-HF and SphI-HF (both from NEB). The digested insert was cloned into the pDVWSK601 plasmid, previously digested with BsrGI-HF and SphI-HF (NEB). The integrity of pDVWSK601 C1218T was verified by full-plasmid sequencing (Microsynth Seqlab).

### *In vitro* transcription of DENV gRNA

DENV2 NGC full length *in vitro* transcripts (IVT) were generated using pDVWSK601, pDVWSK601-LucUbi, and pDVWSK601 C1218T linearized by restriction using XbaI (NEB). Three micrograms of linearized plasmid were incubated at 37°C overnight in a transcription reaction containing 140 U T7 RNA polymerase (Promega), rNTP-Mix (containing 3.125 mM ATP, CTP, and UTP and 1.56 mM GTP, all from Roche), 100 U RNAsin ribonuclease inhibitor (Promega) and 1 mM m^7^G(5’)ppp(5’)G RNA cap analogue (NEB) in 1x RRL buffer (80 mM HEPES-KOH pH7,5, 12 mM MgCl_2_, 2 mM spermidine, 40 mM DTT in HPLC water). RNA was extracted using 0.3M sodium acetate pH 4.5 and acid phenol:chloroform:isoamyl alcohol pH 4.5 (Ambion), followed by isopropanol precipitation and resuspension in nuclease-free water. RNA integrity was determined by using non-denaturing agarose gel electrophoresis and optical density 260/280 nm ratio.

### Virus production and measurement of infectious titers

DENV2 NGC and DENV2 NGC C1218U viral stocks were produced as described elsewhere (28). In brief, 6×10^6^ BHK-21 cells were electroporated with 10 µg of DENV gRNA IVT resuspended in Cytomix solution (29) supplemented with 2 mM ATP and 5 mM glutathione. Each electroporation was conducted at 975 µF and 270 V. Cells were then maintained in 10-cm dish with culture medium supplemented with 15 mM HEPES (Gibco). Virus supernatant was harvested at day 3, 4, 5 and 6 post-electroporation and concentrated with centrifugal filter devices (Centricon plus-70 centrifugal filter, MWCO 100 kDa, Millipore), aliquoted and stored at −80°C. Viral stock infectious titers were determined after one freeze-and-thaw cycle by limiting dilution assay (TCID_50_) using Huh7 cells. In brief, cells were fixed with 4% PFA in PBS 72h post infection and DENV-positive cells stained using a pan-flavivirus antibody produced from the HB112 hybridoma cell line (ATCC) (30).

For DENV2 NGC C1218U viral stocks, RNA was extracted from the supernatant collected daily. Three µg of total RNA were reverse transcribed using the High-Capacity cDNA Reverse Transcription Kit (Applied Biosystems) as recommended by the manufacturer. The region encompassing the C/U substitution was amplified by PCR using the Platinum II Taq Hot-Start DNA Polymerase (Invitrogen) and the following primers: 5’-AGAGAAACCGCGTGTCGACTGTACAACAGCTGACAAAGAG-3’ and 5’-GTGTCATTTCCGACTGCATGCATGCTCTTCCCCTGAGTG-3’. PCR products were purified by gel-extraction (Macherey-Nagel) and subjected to Sanger sequencing (Microsynth Seqlab) to ensure that reversion had not occurred during virus amplification.

### ViREn: a two-step purification and enrichment method for viral RNAs

#### Extraction of DENV RNA extraction from infected cells and viral particles

3×10^6^ Huh7 cells were infected with DENV2 NGC at a multiplicity of infection (MOI) of 10 TCID_50_/ml for 48 h. For the purification of DENV gRNA and sfRNA from infected cells, total RNA was extracted using TRIzol reagent as recommended by the manufacturer (Invitrogen). For the purification of DENV gRNA from viral particles, 15 ml of the culture supernatant was concentrated using centrifugal filter devices (Amicon Ultra Centrifugal Filter 100 kDa MWCO, Milipore) before extraction with TRIzol LS reagent as recommended by the manufacturer (Invitrogen). The resulting RNA pellets were subjected to a stepwise enhanced deproteination and purification method. First, the RNA pellets were resuspended in ribosome disassembly buffer (20 mM Tris-HCl, 0.25 mM MgCl_2_, 150 mM NaCl, 1% Triton X-100, 100 μg/ml cycloheximide, 40 units RNAsin ribonuclease inhibitor (Promega), 90 mM EDTA pH8.0 in DEPC-treated H_2_O) and tumbled at 4°C for 10 min before extraction with TRIzol LS reagent. Next, the RNA pellets were resuspended in proteinase K buffer (20 mM Tris-Cl pH 7.5, 0.5% SDS, 1 mM EDTA in DEPC H_2_O) supplemented with 300 µg/ml proteinase K (Macherey Nagel) and incubated at 42°C for 20 min. Finally, the RNA was extracted using 0.3M sodium acetate pH 4.5 and acid phenol:chloroform:isoamyl alcohol pH 4.5 (Ambion), followed by isopropanol precipitation. RNA was resuspended in nuclease-free H_2_0.

#### Viral RNA separation by sucrose density gradient ultracentrifugation

Linear sucrose gradients were prepared as follows. Sucrose solutions (790 µl) were layered on top of each other starting with 20% (w/v) sucrose solution at the bottom of the tube (4 mL tubes, Beckman Coulter), 16.25% (w/v), 12.5% (w/v), 8.75% (w/v), to 5% (w/v) sucrose in 1x gradient buffer (50 mM KCl, 10 mM potassium acetate, 0.1 mM EDTA, 200 units/ml of RNAsin ribonuclease inhibitor, pH 5.2 in DEPC H_2_O). Each sucrose solution layer was frozen at −80°C for 15 min prior to addition of the next layer. Gradient tubes were thawed overnight in a cold room prior to ultracentrifugation. RNA samples were layered on top of the gradient and ultracentrifuged using a SW60TI rotor (Beckman Coulter) at 43,800 rpm for 4 h at 4°C. Following centrifugation, gradients were manually fractionated from the top (11 fractions, 375 µl each). RNA was extracted from each fraction using 1:1 ratio of extraction buffer (50 mM KCl, 10 mM potassium acetate, 0.1 mM EDTA pH 5.2 in DEPC-treated H_2_O). Samples were heated at 65°C for 10 min after addition of 30 µg of Glycoblue (Invitrogen), 0.3M sodium acetate pH 4.5, and acid phenol:chloroform:isoamyl alcohol pH 4.5 (Ambion). RNA extraction was followed by isopropanol precipitation.

Full-length DENV gRNA IVT that served as negative control for the detection of RNA modifications by BSSeq and DRS was prepared following the same protocol. 100 µg of DENV gRNA IVT were mixed with total RNA prepared from uninfected Huh7 cells and subjected to sucrose density gradient ultracentrifugation and fractionation.

#### Specific affinity capture of DENV RNAs

RNAs extracted from fractions 3 and 11 were subjected to DENV2 gRNA affinity capture using RNA seq MagIC beads - dengue virus - DENV2 - kit (ElementZero Biolabs), containing a pool of 20 different DNA probes binding across DENV gRNA, following manufacturer instructions. Custom beads were generated for DENV2 sfRNA. In short, 20 pmol of beads were used to capture either DENV gRNA or sfRNA from 1 – 3 µg of total RNA. The captured RNA was eluted in nuclease-free water and probes denatured at 92°C.

### Quantitative analysis of DENV RNA by qRT-PCR

Taqman probe-based RT-qPCR was employed to quantify DENV RNAs after purification. Briefly, 3 μl of extracted RNA were mixed with 12 μl qPCR mix (qPCRBIO Probe 1-Step Go Lo-ROX, PCR Biosystems) according to the manufacturer instructions and using the following cycling protocol: 50°C 10 min, 95 °C 1 min, 95°C 10 sec, 60°C 1 min; steps 3 – 4 repeated for 39 times. Absolute RNA copy numbers of DENV were calculated using a standard curve of purified DENV IVT. DENV RNA (gRNA and sfRNA) and cellular housekeeping transcripts (GAPDH, 18S rRNA, and 28s rRNA) were analyzed in triplicates using a CFX96 Touch Real-Time PCR detection system (Bio-Rad). Specific primers and probes are listed in Table 1. Note that a DENV 3’UTR probe was employed for the detection of both DENV2 NGC gRNA and sfRNA and a DENV E probe to detect specifically DENV2 NGC gRNA.

**Table 1:**
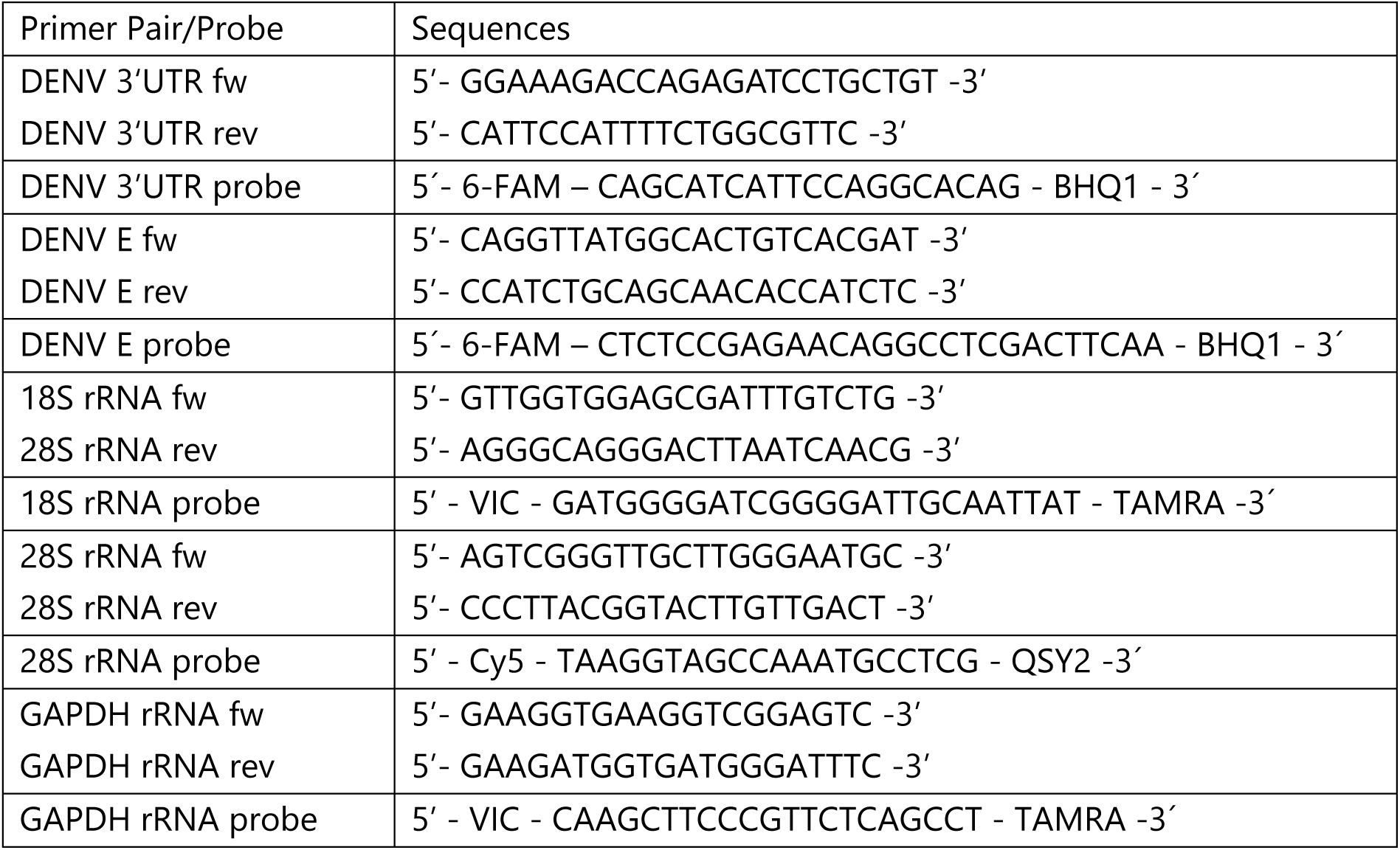
List of primers and probes used for qRT-PCR.

### Sedimentation analysis of DENV RNA by northern blot analysis

RNA from each sucrose fraction was loaded and separated by electrophoresis in a 1% agarose denaturing gel at 110V for 30 min. RNA was then transferred by capillarity overnight to a positively charged nylon membrane (Roche) using 20x SSC buffer (3M NaCl, 0.3M sodium citrate, pH 7.0 in DEPC-treated H_2_O). DENV specific probes, targeting NS1-encoding or 3’ UTR regions were generated freshly using the above-described *in vitro* transcription protocol and DIG-labelled rNTP (Roche) (Table 2). DENV RNA was detected using DIG northern reagents (Roche), following manufacturer instructions. Signals on northern blot membranes were imaged by ECL Chemocam Imager (Intas Science Imaging Instruments). Bands were quantified using LabImage 1D Software (Intas Science Imaging).

**Table 2:**
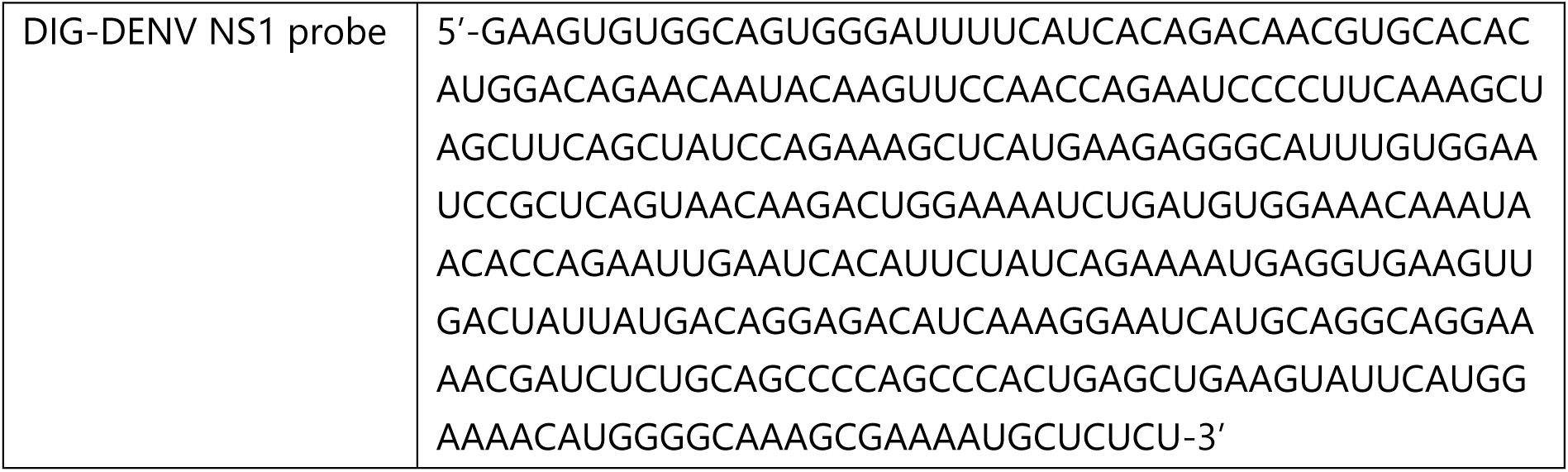
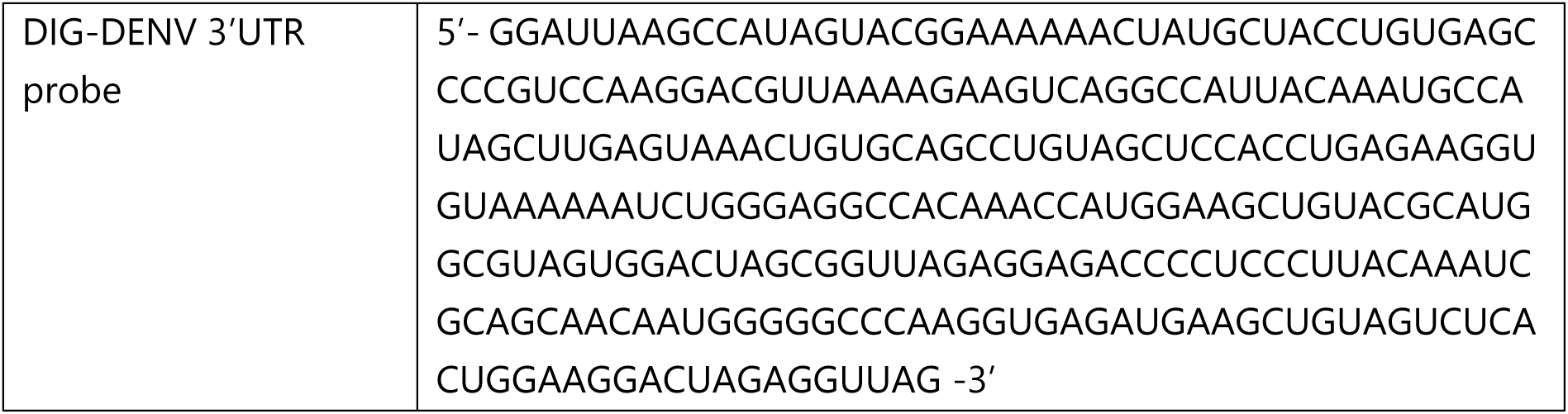
Sequences of DIG-labelled probes used for northern blot.

### Estimation of DENV RNA enrichment after affinity capture by RNA-Seq

Sequencing libraries were produced with the NEBNext Ultra II Directional RNA Library Prep Kit and sequenced on a NextSeq550 System (Illumina). The 80 nt long single-end reads were first aligned to human rRNA sequences as downloaded from the UCSC Genome Browser using bowtie v1.2.2 (31). Reads that did not align to rRNA were further aligned to the human genome (hg38) using STAR v2.5.4b (32), providing a gtf file with the exonic coordinates of the basic set of Gencode V38. Read counts were summarized at the gene level using featureCounts (33), only counting reads that are fully contained within exonic regions. Finally, reads that did not align to human genomic sequences were aligned to the DENV2 NGC full plasmid sequence with bowtie. In all alignment steps, up to two mismatches were allowed. Read coverages on the DENV2 sequence were visualized with IGV (34).

### Illumina-based bisulfite sequencing (BSSeq)

200 ng of DENV gRNA from Huh7 cells or 200 ng DENV full length IVT were used for bisulfite conversion according to Dai and colleagues (35). After bisulfite conversion and desulphonation, RNAs were first end-repaired and purified before being converted to libraries using NEBNext small RNA library kit (NEB). The DNA libraries were quantified using a fluorometer (Qubit 3.0 fluorometer, Invitrogen) and qualified using Agilent TapeStation 4150. Libraries were multiplexed and subjected to high-throughput sequencing on an Illumina NextSeq2000 instrument with a 75 bp single-end read mode.

### Targeted MiSeq bisulfite sequencing

One µg of RNA extracted from fraction 11 or from total RNA was used for bisulfite conversion using the EZ RNA Methylation Kit (Zymo Research). RNA was reverse transcribed to cDNA using High-Capacity cDNA Reverse Transcription Kit as recommended by the manufacturer (Applied Biosystems). The DENV gRNA targeted region (nt 1106-1266), encompassing the CUCCA motif, was amplified by PCR using the Platinum II Taq Hot-Start DNA Polymerase (Invitrogen) using the following primers: forward: 5’-GGAAGTATTGTATAGAGGTAAAG-3’ and reverse: 5’-CCAAATAATCCACATCCATTTCCCC-3’. PCR products were purified by gel-extraction (Macherey-Nagel) and subjected to MiSeq (Illumina) or to sanger sequencing (Microsynth Seqlab). Individual PCR products we barcoded using Nextera XT Index Kit v2 Set A and subjected to MiSeq using MiSeq Reagent Nano Kit v2 (300-cycles Illumina). The occurrence of m^5^C modifications was analyzed and visualized using the web-based pipeline BisAMP (https://bisamp.dkfz.de/index.php?site=4) (36).

### Nanopore direct RNA sequencing (DRS) of DENV gRNA and IVT

Complementary DNA oligos were designed to dissolve DENV gRNA secondary structures by tiling along the RNA (37). In total 100 oligos (Supplementary Table **S1**) were dissolved at 100 µM and mixed equimolar. For DRS, the oligo mix was diluted 1:10. 1 µl oligo mix was combined with 8 µl purified, polyadenylated DENV gRNA or IVT and 1 µl 10x hybridization buffer (f.c. 10 mM Tris pH 7.5, 50 mM NaCl). In a PCR thermocycler, the DENV gRNA was first denatured (5 min, 95°C) and then annealed to the oligos by slowly decreasing the temperature (0.1°C/sec to 22°C). The oligo-annealed DENV gRNA was immediately subjected to DRS library preparation with SQK-RNA004 (ONT) by addition of 3 µl 5x Quick ligation buffer (NEB), 1 µl RTA (ONT) and 1 µl T4 DNA ligase, high concentration (NEB). Adapter ligation was carried out 10 min at room temperature. To resolve remaining secondary structures, a complementary DNA strand was synthesized with Induro Reverse Transcriptase (RT, NEB). To 15 µl ligation, 14 µl H_2_O, 8 µl 5x Induro RT reaction buffer, 2 µl 10 mM dNTPs (Thermo Fisher Scientific) and 1 µl Induro RT were added and incubated 30 min at 60°C, followed by 10 min denaturation at 70°C. Reactions were cleaned up with 1.8x RNA Clean XP beads (Beckman Coulter), washed two times with 80% ethanol and eluted in 13 µl H_2_O. In a total volume of 20 µl, the RLA adapter (ONT) was ligated as described above. Reactions were cleaned up with 1x RNA Clean XP beads, washed two times with ONT wash buffer and eluted in 33 µl REB (ONT). The concentration of the library was determined with the Qubit dsDNA HS kit (Thermo Fisher Scientific). Libraries were loaded on RP4 Promethion flow cells as described in the manufacturers protocol. Sequencing was performed overnight on a P24 A series device equipped with MinKNOW 24.06.14. POD5 files were called with Dorado v0.8.2 and the rna004_130bps_sup@v5.1.0 model and the modified bases option (“--modified-bases m5C inosine_m6A pseU”) enabled. Data were aligned with minimap2 v2.28 and modkit v0.4.2 was used to generate the corresponding bedmethyl files (modification threshold = 0.95, Supplementary Table **S2**).

### Preparation of oligonucleotides for accurate m^5^C Nanopore reads basecalling

#### RNA oligonucleotide phosphorylation and ligation

Sequencing reads from m^5^C-modified and unmodified oligonucleotides were generated to train ONT’s dorado basecaller. Unmodified 5’ and 3’ oligonucleotides were purchased from biomers.net. The central m^5^C-modified oligonucleotide was purchased from Dharmacon (see table 3). An equimolar mixture of the three RNA oligonucleotides with modified or unmodified central oligonucleotide) were phosphorylated at the 5’ end using 0.75 U/μl T4 polynucleotide kinase (Thermo Fisher Scientific) per 30 μM of RNA in 1x kinase-ligase buffer (50 mM Tris-Cl pH 7.4, 10 mM MgCl₂, 5 mM DTT, 2 mM ATP) for 60 min at 37°C. The phosphorylated oligonucleotides were immediately used for ligation by adding 2% less complementary DNA splint to the reaction mixture. For this experiment, RNA oligonucleotides of 900 pmol each were hybridized to 882 pmol of DNA splint by heating at 75°C for 4 min, followed by slow cooling to room temperature over 15 min.

**Table 3:**
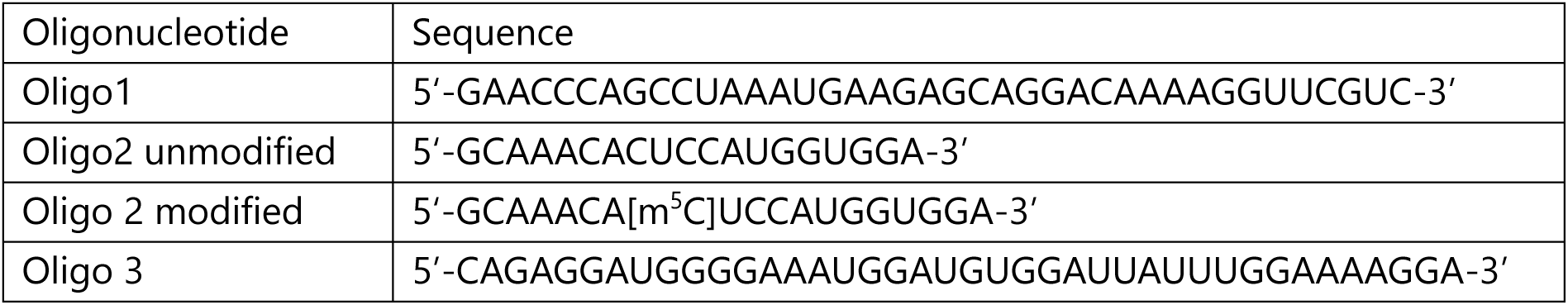
Sequences of RNA oligonucleotides used for ligation.

Ligation was initiated by adding 2 U/μl T4 DNA Ligase (Thermo Fisher Scientific), and the reaction was incubated overnight at 16°C. To remove the DNA splint, DNase I (Thermo Fisher Scientific) was added at a final concentration of 0.1 U/μl, followed by incubation at 37°C for 30 minutes. The ligated construct was purified from unligated oligonucleotides using a 8% denaturing polyacrylamide gel and electrophoresis. RNA purity and concentration were assessed using a NanoDrop 2000 (Thermo Fisher Scientific).

#### Polyadenylation and DRS library preparation

The ligated RNA m^5^C-modified and unmodified oligonucleotides were polyadenylated at the 3’ end using E. coli poly(A) polymerase (NEB) according to the manufacturer protocol. The RNA was then purified using the Oligo Clean & Concentrator Kit (Zymo Research) before proceeding to library preparation. Library preparation was performed using 1 µg of polyadenylated RNA with the SQK-RNA004 kit (ONT), following the manufacturer standardized protocol. In this step, SuperScript III Reverse Transcriptase (Invitrogen) was used instead of Induro Reverse Transcriptase (New England Biolabs). The prepared libraries were quantified using the Qubit dsDNA HS Assay Kit (Invitrogen), loaded onto MinION flow cells, and sequenced with MinKNOW (Version 24.02.26) for 72 h.

### Analysis of NSUN6 expression during DENV infection by western blot analysis

Huh7 cells (1×10^5^ cells/well) were seeded into a six-well dish one day before infection with DENV2 NGC at an MOI of 10 or mock treatment. Medium was replaced 6 h after the addition of the virus and cells were harvested at the indicated time points after virus addition. Cells were washed with PBS and collected by scraping into ice-cold protein lysis buffer (50 mM Tris-HCl pH 7.4, 150 mM NaCl, 1% Triton X-100, supplemented with 60 mM β-glycerophosphate, 15 mM 4-nitrophenylphosphate, 1 mM sodium orthovanadate, 1 mM sodium fluoride, and EDTA-free cOmplete protease inhibitor cocktail (Roche)). Cells were lysed for 30 min on ice followed by centrifugation for 15 min at 16,000×g at 4°C. Total protein concentration of clarified lysate was determined by the Bradford method using Protein Assay Dye Reagent (Bio-Rad) and absorbance measurement at 595 nm.

Equal amounts of total protein (20 µg) were denatured for 5 min at 95°C in reducing 1× Laemmli buffer (62.5 mM Tris-HCl pH 6.8, 8.33% glycerol, 1.5% SDS, 1.5% β-mercaptoethanol, 0.005% bromophenol blue) and separated on a 10% resolving gel by SDS-polyacrylamide gel electrophoresis in buffer containing 25 mM Tris-HCl pH 8.8, 192 mM glycine, and 0.1% (w/v) SDS. Proteins were transferred to a 0.45 µm polyvinylidene difluoride membrane (Millipore) by tank blotting at 360 mA for 2.5 h at 4°C in transfer buffer (25 mM Tris-HCl pH 8.3, 150 mM glycine, and 20% (v/v) methanol). Membranes were blocked with 5% (w/v) powdered milk (Roth) in Tris-buffered saline containing 0.1% Tween 20 (50 mM Tris-HCl pH 7.4, 150 mM NaCl, 0.1% Tween 20) for 1 h. Immunostaining was performed in blocking buffer by incubating membrane sections overnight at 4°C with rabbit polyclonal anti-NSUN6 (GeneTex, 1:1,000), mouse monoclonal anti-GAPDH (Santa Cruz Biotechnology, 1:5,000), or rabbit polyclonal anti-DENV NS5 (GeneTex, 1:1,000), followed by incubation for 2 h with goat polyclonal secondary antibodies conjugated to horseradish peroxidase (all from Sigma-Aldrich). Proteins were detected by chemiluminescence using Western Lightning Plus enhanced chemiluminescence (ECL) reagent (Perkin Elmer) and detection on an ECL Chemocam Imager (Intas Science Imaging). Band intensities were quantified using Image Studio Lite software (Li-Cor, version 5.2.2). The statistical comparison between DENV infected vs Mock samples were performed using by two-way ANOVA with multiple comparison (n.s., non-significant, GraphPad Prism v8.4.3).

### Generation of Huh7 NSUN6 knockout cell clones by CRISPR/Cas9

Huh7 knockout (KO) cell clones were generated using a two-CRISPR RNA (crRNA) approach as described in (38). Two crRNAs targeting the NSUN6 locus at exon 1 and 3 (5’-/AltR1/GCATTTCTCTTGGTGGAGAAC/AltR2/-3’, 5’-/AltR1/CATGACAGAGTCTAAAGTCTGC/AltR2/-3’) were purchased from IDT. In brief, non-targeting (NT) control cell lines were generated with the CRISPR-Cas9 negative control crRNA#1 (IDT). Individual crRNAs and tracrRNA (IDT) were resuspended TE buffer (10 mM Tris, 0.1 mM EDTA, pH 7.5) to a final concentration of 200 µM and hybridized at equimolar amounts by heating to 95 °C for 5 min and slow stepwise cooling to room temperature. Pre-assembled crRNA/Cas9 ribonucleoprotein complexes (RNPs) were obtained by gently mixing 1.2 µl of hybridized RNAs with 17 µg of recombinant Cas9 protein (IDT) and 2.1 µl of PBS, followed by 20 min incubation at room temperature.

2×10^6^ Huh7 cells were resuspended in 91 µl supplemented Solution T (Lonza) and mixed with 2.5 µl of each pre-assembled RNP and 4 µl electroporation enhancer (IDT). Nucleofection was performed using an Amaxa 2b nucleofector (Lonza) and pre-set program T-22. Nucleofected cells were mixed with pre-warmed conditioned medium and seeded. After 48 h cells were seeded for clonal amplification in 96-well plates. Homozygous clones were screened using target-specific PCR (Platinum II Taq hot-start DNA polymerase, Invitrogen) on genomic DNA using the following primers: NSUN6-KO-For: 5’– TCGGCTAGGCTTGAGAAA GC –3’ and NSUN6-KO-Rev: 5’– AGGGTTTCGCCATACTGG –3’. The expression of NSUN6 in homozygous cell clones was further analyzed by immunoblotting.

For further analysis, Huh7 cell pools were prepared to account for single cell clone variation. Equal numbers of Huh7 NT cell clones (#1, 2, 3, 4 and 8) and Huh7 NSUN6 KO cell clones (#1, 19, 29 and 33) were mixed and frozen. Cell pools were thawed and used for experiments within 5 passages.

### Characterization of NSUN6 KO cell clones

#### Analysis of cell growth

Cell growth was measured using IncuCyte live-cell analysis system (Sartorius). 3×10^3^ cells per well were seeded in a 96-well plate. Cells were incubated at 37°C for overnight before starting the live-cell analysis for 5 days. Specifically, cell growth was measured by imaging 4 fields of view for each well at 3-h interval. Cells were counted, and growth curves generated using the IncuCyte live-cell analysis system. Values were normalized to time 0. Statistical analysis was performed using repeated measures two-way ANOVA with Geisser-Greenhouse’s correction (n.s., non-significant; ∗, p < 0.05; ∗∗, p < 0.01; ∗∗∗, p < 0.001; GraphPad Prism v8.4.3).

#### Measurement of translation

The global translation of the Huh7 NT and NSUN6 KO cells was measured using Click-iT Plus OPP Alexa Fluor 488 Protein Synthesis Assay Kit (Invitrogen) and flow cytometry (BD FACSCelesta Cell Analyzer). 2×10^5^cells was seeded in triplicates in 6-well plates the day before measurements. Huh7 NT cells treated with 200 µg/ml cycloheximide for 1 h were used as negative control. Cells were treated with 50 µM OP-puro for 1 h at 37°C before fixation with 4% PFA in PBS on ice (in the dark) for 15 min. Cells were washed once with 2 ml PBS before membrane permeabilization with 1 ml 3% FBS, 0.1% saponine in PBS for 5 min. Cells were resuspended in Click-it reaction buffer as recommended by the manufacturer and washed twice with PBS supplemented with 3% FBS and 0.1% saponine before analysis by flow cytometry. Statistical analysis of the technical replicates was performed using unpaired t-test (n.s., non-significant; ∗, p < 0.05; ∗∗, p < 0.01; ∗∗∗, p < 0.001; GraphPad Prism v8.4.3).

#### LC-MS/MS analysis of tRNA^Thr^ m5C levels in Huh7 NSUN6 KO cells

Purification of tRNA^Thr^ was based on a method by Kristen and colleagues (39). In detail, 6 µg of purified total tRNAs from either Huh7 NT or NSUN6 KO cells were mixed with 30 pmol of each FAM-labelled cDOQ (see table 4) (all purchased from IDT) in a hybridization buffer (30 mM HEPES pH 7.5, 100 mM potassium acetate). Hybridization was performed by incubating at 95 °C for 1 min, followed by a stepwise reduction (5 °C/30 s) up to 70 °C, followed by 5 °C/min to 30 °C, and 2.5°C/min to 20 °C. Samples were loaded on a 10% native polyacrylamide gel prepared according to supplier’s protocol (Carl Roth). Gel was run for 1:30 h at 45 mA followed by visualization using a Typhoon Trio+ (ex: 488 nm, em: 520 nm BP, Amersham Biosciences). Shifted bands were cut with a scalpel, smashed and resuspendend in 500 µl 500 mM ammonium acetate (Carl Roth). Gel pieces were frozen at −80 °C, then incubated at 15 °C overnight under slight shaking. The next day, solutions were filtered using 0.45 µm Nanoseps (PALL) and precipitated using 1 µl GlycoBlue (Invitrogen) and 1 ml ethanol. Afterwards, DNA oligonucleotides were digested using 2 units DNase I-XT (NEB) in its respective reaction buffer for 1 h at 37 °C. Finally, samples were purified using RNA Clean & Concentrator-5 (Zymo Research) and eluted in 6 µl nuclease-free water.

**Table 4:**
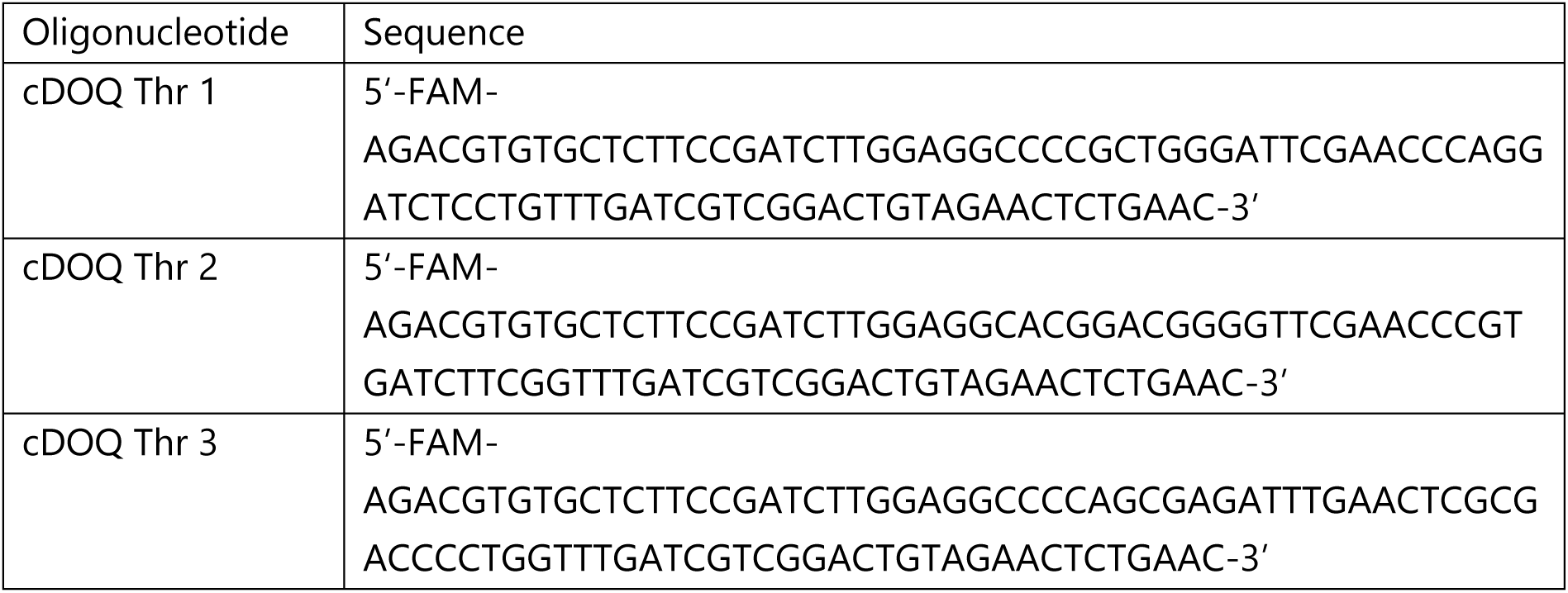
Sequences of oligonucleotides used for DORQ-seq.

200 ng of tRNA^Thr^ purified from Huh7 NT or NUSUN6 KO cells were digested with 0.2 U snake venom phosphordiesterase from *C. adamanteus* (Worthington), 10 U benzonase (Sigma-Aldrich), 0.2 U bovine intestine phosphatase (Sigma-Aldrich), 0.6 U nuclease P1 from *P. citrinum* (Sigma-Aldrich) and 200 ng pentostatin (Sigma-Aldrich) in a total volume of 20 µl buffer (5 mM Tris pH 8, 1 mM MgCl_2_). Reaction was incubated for 2 h at 37 °C and stopped by addition of 30 µl nuclease-free water. The LC-MS/MS device was an Agilent 1260 Infinity (II) with an Agilent 6460A triple quadrupole mass spectrometer equipped with an electrospray ion source with following parameters: gas temperature 350 °C, gas flow 8 L/min, nebulizer pressure 50 psi, sheath gas temperature 350 °C, sheath gas flow 12 L/min, capillary voltage 3000 V and nozzle voltage 0 V. A Synergi 4 µm Fusion-RP 80 Å 250 x 2 mm (Phenomenex) was used as column. Separation was achieved using a two-component gradient with solvent A (5 mM ammonium acetate pH 5.3 (Sigma-Aldrich), adjusted with acetic acid, supplemented with 1% (v/v) acetonitrile (Honeywell)) and solvent B (acetonitrile with 1% (v/v) nuclease-free water). The gradient started with 100% solvent A, followed by a linear increase to 8% solvent B within 10 min. Solvent B was further increased to 40% within another 10 min to finally decrease to 100% solvent A in 3 min. Subsequent equilibration was performed for 7 min. Flow rate was constant at 0.35 mL/min. Canonical nucleosides were detected using a diode array detector at 254 nm while modified nucleosides were detected using dynamic multiple reaction monitoring (dMRM) in positive mode with following parameters: m^5^C mass transition: 258 -> 126.1, retention time: 7.9 min; m^3^C mass transition: 258 -> 126, retention time: 4.4 min. Quantification was performed as previously described (40). In short, external calibration was performed by measuring increasing amounts of modified nucleoside standards (1 to 5000 fmol) or canonical nucleoside standards (0.5 to 500 pmol). Internal calibration was performed by addition of ^13^C-labeled internal standard from ^13^C-fed yeast. The respective slopes of the calibration curves were used to calculate the amounts. Modified nucleosides were set in relation to the detected adenosine amount of the respective sample. Results were set in relation to NT.

### Measurements of viral replication

Viral replication was measured using DENV2 NGC *firefly* luciferase reporter RNA as previously described (28). Five μg of DENV full-length *firefly* reporter IVT were electroporated into 1×10^6^ of Huh7 NT or NSUN6 KO cells suspended in Cytomix solution completed with ATP and glutathione. Each electroporation was conducted at 500 μF and 166 V. For the time points 4 h and 24 h post-electroporation, 1.1×10^5^ were then seeded in duplicates in a 24-well plate. For the time points 48 h, 72 h and 96 h, 6.5×10^4^ cells were seeded in duplicates in a 24-well plate. Half of the cells were co-treated with 27 µM NITD008 (Tocris Bioscience) to inhibit DENV RNA-dependent RNA polymerase NS5 activity and viral RNA synthesis (41). Cells were harvested at the indicated time points by addition of luciferase lysis buffer (25 mM Gly-Gly pH 7.8, 15 mM MgSO_4_, 15 mM K_3_PO_4_, 4 mM EGTA, 10% glycerol, 0.1% Triton X-100 supplemented with 1 mM DTT) and immediately stored at −80°C. For measurements of the *firefly* luciferase activity, 100 µg cell lysates were mixed with 360 µl assay buffer (25 mM Gly-Gly pH 7.8, 15 mM MgSO_4_, 15 mM K_3_PO_4_, 4 mM EGTA) supplemented with 1 mM DTT, 2 mM ATP and 70 µM D-luciferin (P.J.K). All measurements were done in duplicate using a tube luminometer (Berthold Technologies). Replication efficiency was determined by normalization to the 4 h values, which reflects transfection efficiency. Statistical analysis was performed using repeated measures two-way ANOVA with Geisser-Greenhouse’s correction (n.s., non-significant; ∗, p < 0.05; ∗∗, p < 0.01; ∗∗∗, p < 0.001; GraphPad Prism v8.4.3).

### NSUN6 *in vitro* methylation assay

#### Bacterial expression and purification of human full-length NSUN6

Plasmid used for overexpression is available at Addgene (ID#188060). Transformed *E. coli* Rosetta (DE3) pLysS cells were grown in terrific broth medium at 37°C till optical density at 600 nm wavelength reached 0,7. After precooling to 16°C overexpression was induced by 500 µM isopropyl-β-D-thiogalactopyranosid (IPTG) for 18-20 hours. Cell pellets were flash frozen in liquid nitrogen and stored at −80°C. Cell pellets were disrupted in lysis buffer (50 mM sodium phosphate pH 7.5, 150 mM NaCl, 10 mM imidazole, 0.1% polysorbate-20). Ni affinity chromatography was performed with HisTrap HP 5 ml (Cytiva) column. Protein was eluted with the elution buffer (50 mM sodium phosphate pH 7.5, 150 mM NaCl, 500 mM imidazole, 0.1% polysorbate-20) and was further purified with the size exclusion chromatography on the Superdex 16/600 75 pg column (Cytiva), equilibrated with SEC buffer (25 mM HEPES pH 7.5, 300 mM NaCl, 1 mM DTT, 0.1% polysorbate-20, 10% glycerol). NSUN6 protein was concentrated with the help of Amicon Ultra Centrifugal Filter (10 kDa, MWCO, Millipore), then aliquoted, flash frozen with liquid nitrogen, and kept at −80°C until further use. The purity and quantity of the recombinant proteins were assessed by SDS-PAGE followed by staining with Coomassie blue.

#### Tritium incorporation assay for NSUN6

RNA methylation was carried out in triplicates for 60 min in the reaction buffer (50 mM TRIS-HCl pH7, 50 mM NaCl, 5mM MgCl_2_, 1mM DTT, 5% DMSO). Recombinant NSUN6 was added at the final concentration of 30 nM, together with the methyl group donor S-adenosyl-L-methionine (SAM). A mixture of cold SAM (NEB) and ^3^H-SAM (Hartmann Analytics) was added to the final concentrations of 1.2 µM and 0.038 µCi µl^−1^. Different DENV RNA substrates were added to the same molar concentration of 750 nM. The tRNA^Thr^, a known NSUN6 substrate, was used as a positive control at the same molar concentration and refolded by 5 min incubation at 75°C. An IVT of the eGFP-coding sequence, containing no CUCCA motifs, was used as negative control. 8 µl of solutions were spotted onto the filters (Whatman) and placed into the ice-cold 5% trichloroacetic acid (TCA) (Sigma-Aldrich) for RNA precipitation and incubated for at least 15 min. All other washing steps were performed at room temperature, namely two more in TCA for 20 and 10 min, and the last one in ethanol for 10 min. Washed filters were dried in the fume hood overnight or under the infrared lamp for 40-60 min. Dry filters were transferred into scintillation vials with 3 ml of Ultima Gold liquid scintillation cocktail (Sigma-Aldrich). The relative amount transferred onto RNA 3H-methyl groups was measured as counts per minute (CPM) with the Tri-Carb 2810TR Low Activity Liquid Scintillation Analyzer (Perkin Elmer). Statistical analysis of the technical replicates was performed using unpaired t-test (n.s., non-significant; ∗, p < 0.05; ∗∗, p < 0.01; ∗∗∗, p < 0.001; GraphPad Prism v8.4.3).

### Measurement of DENV gRNA turnover

2×10^5^ cells per well were seeded in 6 well-plates the day before infection. Cells were infected by DENV2 NGC at a MOI of 0.1. Twenty-four hours post infection, cells were treated with 27 µM NITD008 (Tocris Bioscience) to inhibit viral RNA synthesis and with DMSO, as vehicle control. Cells were harvested at 24-h interval for up to 5 days after treatment with NITD008. Total RNA was extracted with TRIzol reagent (Invitrogen) as recommended by the manufacturer. DENV gRNA copies were quantified by Taqman-probe based qRT-PCR as described above. DENV gRNA levels were normalized to that of *GAPDH* mRNA. Fold changes in abundance relative to time point 24 h were calculated using the ΔΔCT method (42) and expressed as a percentage. The resulting decay curves were obtained by using a non-linear regression (GraphPad Prism) to estimate the mRNA half-life (t_1/2_). For half-lives exceeding 100 h or which could not be determined, values are indicated as “n.d. (> 100 h)”. The statistical comparison of the decay curves was performed using by two-way ANOVA (n.s., non-significant; ∗, p < 0.05; ∗∗, p < 0.01; ∗∗∗, p < 0.001; GraphPad Prism v8.4.3).

## RESULTS

### ViREn: a two-step method to purify and enrich DENV RNA

Enrichment and purity of target RNA are crucial for enhancing sequencing coverage and enabling RNA modification mapping, particularly for low-abundance RNAs. We set out to develop a two-step viral RNA purification and enrichment approach – called ViREn – for the isolation of DENV RNA from infected cells and virions (Fig. **1A**). This method is based on the sedimentation coefficient of viral RNAs in sucrose density gradients. Early studies showed the presence of two DENV RNA species in infected cells, which separated in distinct fractions during gradient ultracentrifugation (43,44). The first species corresponded to gRNA, with a sedimentation coefficient of 40-45S. The second, the origin of which was unknown at the time, had a sedimentation coefficient of 6-8S. According to our current knowledge, this second DENV RNA species was likely sfRNA.

**Fig. 1.**
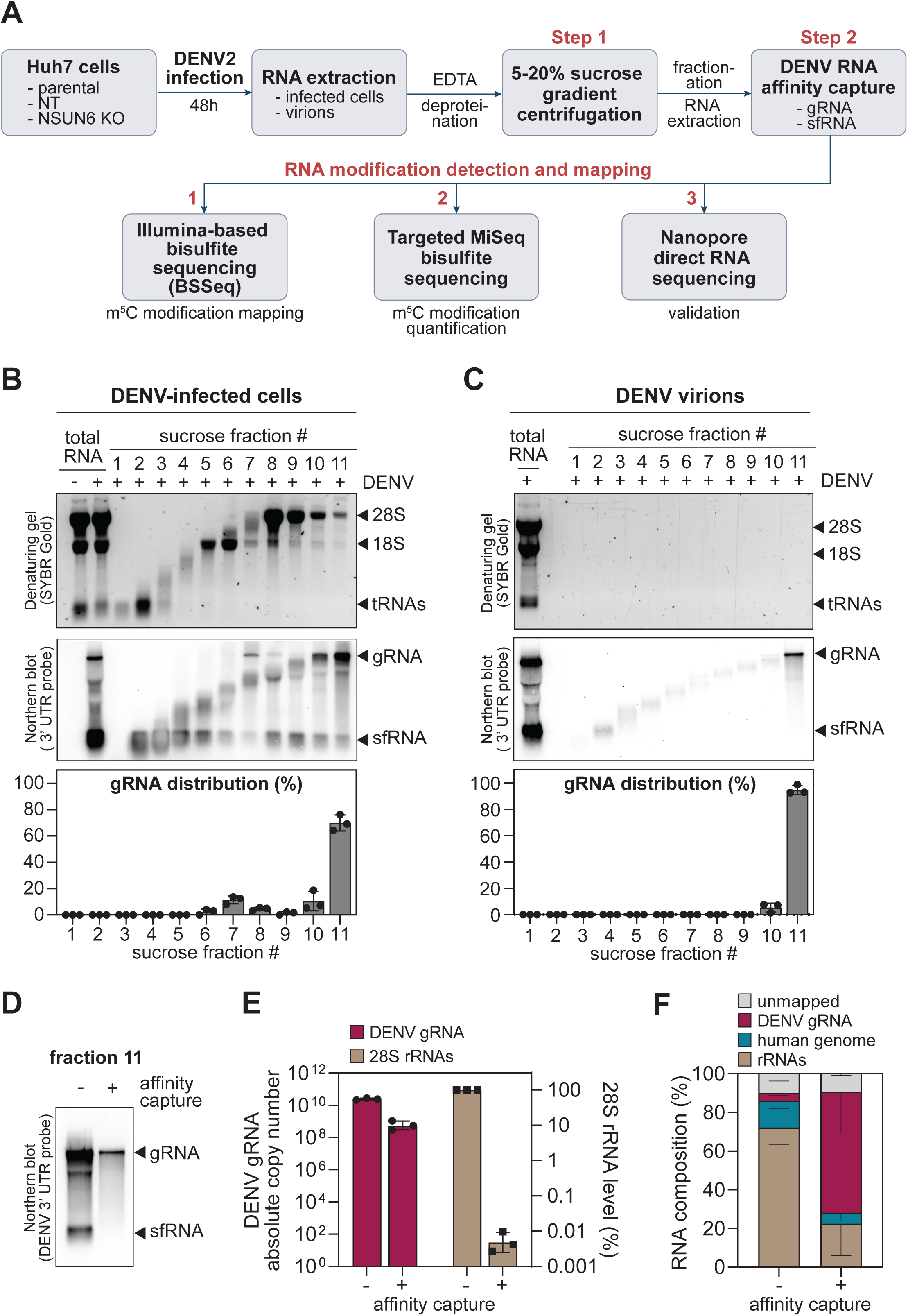
Establishment of ViREn. (**A**) Schematics of ViREn workflow. DENV RNA from infected cells and virions, separated by sucrose gradient ultracentrifugation. DENV gRNA and sfRNA were further enriched using specific affinity capture. The purified DENV gRNA was subjected to RNA sequencing-based approaches for the identification and mapping of m^5^C modifications. (**B** and **C**). RNA extracted from the gradient fractions were analyzed using northern blotting. Shown is a representative analysis (n=3). The distribution of cellular RNAs and DENV RNAs extracted from infected cells (B) and virions (C) across sucrose density gradient fractions was detected using a denaturing agarose gel and corresponding northern blot using a hybridization probe targeting DENV 3’UTR. The quantification of DENV gRNA in each fraction is shown at the bottom (mean ± SD, n=3). (**D**) DENV gRNA was collected in fraction 11 was further purified by affinity capture. Its integrity was visualized by northern blotting. (**E**) The level of DENV gRNA and 28S rRNA contaminant, before and after affinity capture, was measured by Taqman probe-based qRT-PCR (mean ± SD, n=3). (**F**) The total RNA content of fraction 11 before and after affinity capture assay was analyzed by Illumina-based RNA-Seq (n=3). Shown is the proportion of reads in each sample corresponding to DENV gRNA, human genome, rRNAs, and unmapped reads (mean ± SD, n=3)

We first extracted total RNA from Huh7 cells infected with DENV for 48 h. To increase the recovery rate of gRNA, especially of gRNA associated with translating ribosomes, polysomes were disrupted by treatment with EDTA and RNA was extensively deproteinated using proteinase K (Supplementary Fig. **S1A** and **B**). Next, the RNA was separated on a 5-20% sucrose density gradient by ultracentrifugation (Fig. **1A**). The distribution of DENV RNA across the gradient was analyzed after fractionation and RNA extraction, using northern blot and labelling with a probe targeting the DENV 3’ UTR. Consistent with previous reports, the majority of DENV gRNA sedimented in the fraction with the highest sucrose density (fraction 11), separating it from most cellular RNAs (Fig. **1B**). A second band was detected in all gradient fractions, with highest abundance in fraction 3 (Fig. **1B** and Supplementary Fig. **S1C**). This band corresponded to DENV sfRNA, as confirmed by the absence of the signal when using a hybridization probe targeting the NS1-coding sequence at the 5’ end of DENV gRNA (Supplementary Fig. **S1C**). ViREn was applied to gRNA extracted from viral particles. As anticipated, most of the gRNA sedimented in fraction 11 (Fig. **1C**). Importantly, the separation on the gradient effectively removed contaminating viral RNA degradation products, which are likely released into the culture medium as a result of virus-induced cell death (Fig. **1C**).

The second step of ViREn aimed at specifically enriching DENV gRNA and sfRNA by sequence-specific affinity capture, thereby reducing contaminating RNAs that might have co-sedimented. Importantly, this step did not compromise the integrity of the DENV gRNA while removing degraded gRNA and sfRNA (Fig. **1D**). This step of positive selection effectively reduced rRNAs by almost 5 logs as measured by qRT-PCR (Fig. **1E**) and eliminated any remaining sfRNA present in fraction 11 (Fig. **1D**). To better assess the performance of this second step, we performed RNA-Seq analysis on the RNA extracted from fraction 11 before and after affinity capture. On average, the gRNA was enriched by 15-fold after purification (Fig. **1F**). The sfRNA-specific affinity capture proved to be more efficient with a 28-fold enrichment after capture (Supplementary Fig. **S1E** and **F**). Overall, the ViREn method enabled the efficient enrichment and purification of DENV gRNA and sfRNA for further analysis of RNA modifications at single-base resolution.

### Identification of a single m^5^C-modified site at position 1218 of DENV gRNA

We focused our interest on the identification of m^5^C modifications in DENV gRNA because of the availability of several methods, allowing for high-confidence detection and mapping at a single-base resolution (45,46). DENV gRNA purified from cells and from virions was subjected to bisulfite conversion (46,47) coupled to Illumina-based RNA-Seq (BSSeq) (Fig. **2A** and Supplementary Fig. **S2**). Sources of background noise were reduced by (i) using deamination conditions allowing almost complete conversion of C residues to U, and (ii) comparative analysis with an unmethylated *in vitro* transcript (IVT) of the full-length DENV gRNA sequence, which was mixed with total RNA from uninfected cells and subjected to sucrose gradient ultracentrifugation. Analysis was performed in three biological replicates. Stringent analysis of the cytosine pattern revealed a prominent m^5^C site in all three replicates of the gRNA purified from infected cells, at position 1218 in the envelope protein (E) coding sequence (Fig. **2A** and Supplementary Fig. **S2**). Strikingly, this modification was not detectable in gRNA purified from virions, suggesting the potential exclusion of m^5^C-modified gRNAs from packaging into viral particles. The occurrence of m^5^C at position 1218 was confirmed by targeted MiSeq bisulfite sequencing (36,46), which established an average methylation frequency of approximately 20% (Fig. **2B**).

**Fig. 2.**
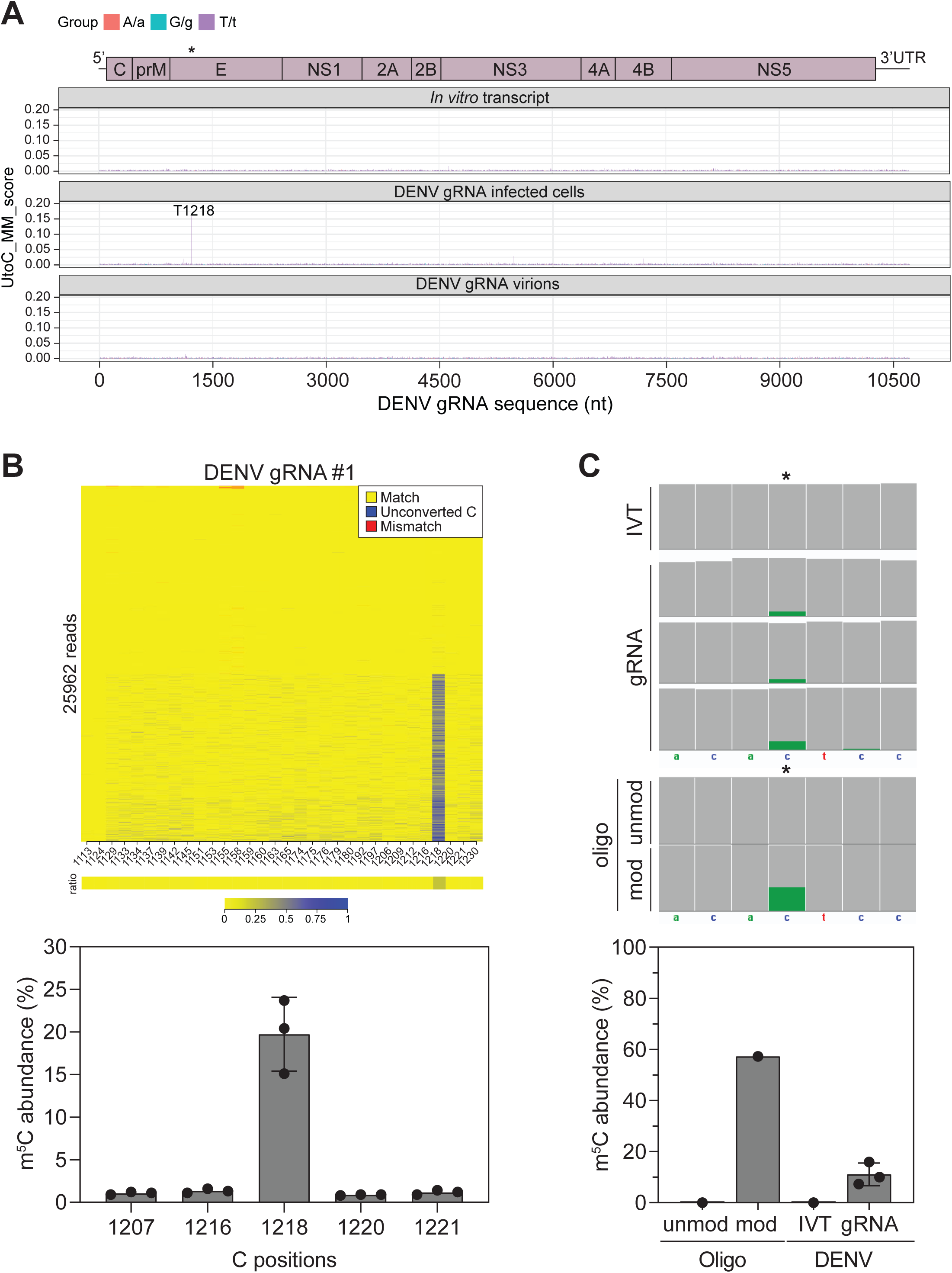
Mapping of m^5^C on DENV gRNA. (**A**) DENV gRNA purified from infected cells and virions were subjected to BSSeq (n=3). Unmethylated *in vitro* transcript processed with the ViREn method was used for comparative analysis and employed to reduce background noise. A representation of DENV gRNA is shown on the top. The asterisk marks the m^5^C position at nt 1218 in the E-coding sequence of DENV gRNA. A schematic of DENV gRNA organization is shown on the top. (**B**) The methylation level of m^5^C1218 detected from infected cell samples was quantified using targeted MiSeq bisulfite sequencing (n=3). Shown is a representative methylation heatmap generated using BisAMP (36). The methylation level of all cytosines in the amplicon (nt 1106-1266) are shown in the panel below. Methylation quantification is shown at the bottom for C1218 and 4 adjacent cytosines (n=3). (**C**) DENV gRNA purified from infected cells was subjected to DRS. (n=3). Unmethylated *in vitro* transcript (IVT) was employed to reduce background noise. Short oligonucleotides either containing m^5^C1218 (oligo mod.) or C1218 (unmod.) were sequenced to assess the accuracy of the modification aware ONT basecaller Dorado. Upper part: IGV coverage snapshots of C1218 and adjacent nucleotides. Reads with a modification probability above 0.95 are labeled green. Lower part: the methylation level of m^5^C1218 was calculated applying a modification probability threshold of 0.95 (mean ± SD, n=3 for DENV gRNA).

To further consolidate this result, we assessed the presence of this modification in native DENV gRNA molecules and established a DENV-specific DRS approach. For this, three substantial challenges had to be overcome: 1) the length of DENV gRNA, 2) potential complex secondary and tertiary structures (48,49), and 3) the lack of a poly(A) tail to pull the gRNA through the nanopore. Therefore, DENV gRNA was polyadenylated at the 3’ end *in vitro*, and complementary DNA oligonucleotides were hybridized along the sequence to disrupt potential RNA structures before passing through the nanopore. To assess the accuracy of ONT’s basecaller Dorado, which was trained to detect m^5^C sites on RNA, we sequenced synthetic unmodified and m^5^C-modified oligonucleotides containing C1218 in its native sequence context (Fig. **2C**). With the stringent probability threshold applied here (see Methods), we underestimated the m^5^C modification levels in the fully m^5^C-modified oligonucleotide but reduced false positive calls to almost zero (Fig. **2C**). In addition, sources of noise were reduced by analyzing three biological replicates of purified DENV gRNA and by comparative analysis with the unmethylated DENV full-length IVT processed alongside. In line with the results obtained by BSSeq, DRS confirmed the presence of a single m^5^C modification at position 1218 of DENV gRNA purified from infected cells, with an occurrence of approximately 10% (Fig. **2C**). This occurrence was lower than that estimated by targeted MiSeq bisulfite sequencing (Fig. **2B**), probably due to the strict thresholding applied for the detection of m^5^C in Dorado.

Altogether, using complementary mapping methods and stringent analysis, we uncovered the presence of a single m^5^C modification at position 1218 of DENV gRNA, at a frequency ranging between 10% and 20%. This occurrence aligns with the median level of m^5^C methylation deposited by m^5^C methyltransferasesin human mRNAs, ranging from 15% to 18% (50,51). Notably, m^5^C1218 is found exclusively in the cytoplasmic gRNA, suggesting a possible sorting process that excludes m^5^C-modified gRNA molecules from packaging into viral particles.

### The host methyltransferase NSUN6 catalyzes DENV gRNA m^5^C1218

The majority of m^5^C modifications on mRNAs is catalyzed by the human methyltransferase NSUN2 (45), whose localization is mainly nuclear (35,50). Among the members of the NSUN family, NSUN6 is a cytosolic methyltransferase that primarily installs m^5^C to the first cytosine of the consensus sequence motif CUCCA, often located in the loop of hairpin structures in mRNA 3’ UTRs (45,52,53) and at the 3’ end of tRNA^Thr^ and tRNA^Cys^ at position 72 (54). Inspection of the DENV gRNA sequence revealed that m^5^C1218 coincided with the first cytosine of a CUCCA motif. Thus, we investigated the potential role of NSUN6 in DENV gRNA methylation, noting that the protein expression of NSUN6 was not altered during infection (Fig. **3A**). We generated Huh7 NSUN6 knockout (KO) single cell clones using the CRISPR/Cas9 technology and two guide RNAs targeting *NSUN6* locus at exon 1 and 3 (Supplementary Fig. **S3A**). Control cell clones were generated with a non-targeting guide RNA (NT) for comparison (Supplementary Fig. **S3B**). To account for the inherent variability of single cell clones, homozygous clones in which the endogenous locus was deleted (Supplementary Fig. **S3A**) and NSUN6 expression was undetectable (Supplementary Fig. **S3B**) were pooled in equal numbers for all subsequent experiments.

**Fig. 3.**
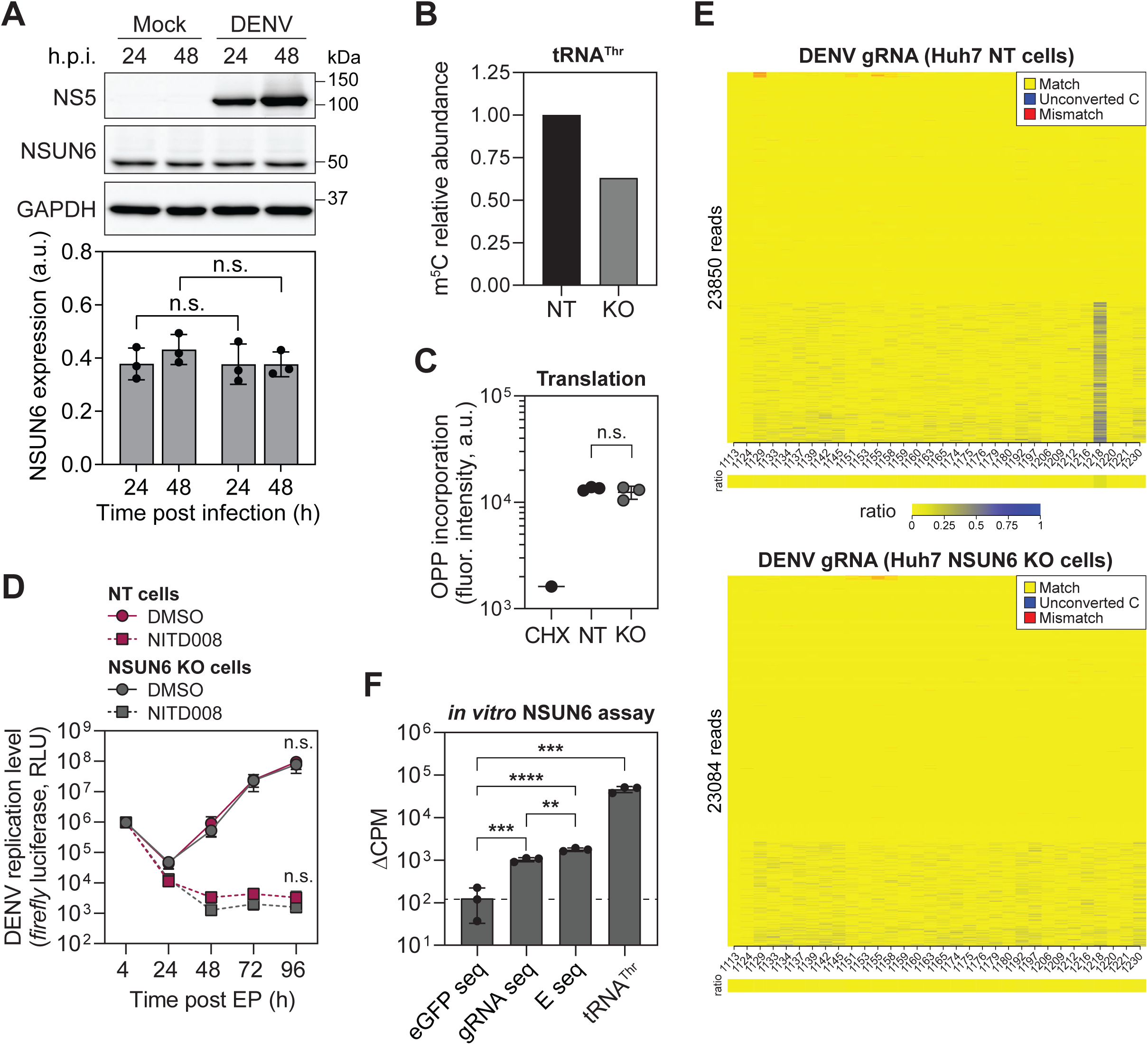
The host methyltransferase NSUN6 catalyzes DENV gRNA m^5^C1218. (**A**) NSUN6 expression levels during DENV infection (n=3). Shown is a representative immunoblot (top) and quantifications. DENV NS5 and GAPDH expression are shown. Levels were normalized to the level of the loading control GAPDH (mean ± SD, n=3). Statistical significance compared to the corresponding mock time point is indicated on the top (n.s., non-significant). (**B**) Analysis of m^5^C levels on tRNA^Thr^ in Huh7 NT and NSUN6 KO cells using the DORQ-seq approach followed by LC-MS/MS (n=1). (**C**) Global translation level of Huh7 NT and NSUN6 KO cells was measured using fluorescently labeled OP-Puro incorporation and flow cytometry (n=1). Statistical significance is indicated on the top (n.s., non-significant). (**D**) DENV full-length *firefly* luciferase reporter was used to measure DENV replication (mean ± SD, n=3). Huh7 NT and NSUN6 KO cells were electroporated (EP) with DENV *firefly* luciferase reporter virus IVT. *Firefly* activity was measured as surrogate for viral replication at 24-h interval for 4 days. Values were normalized to the 4 h time point to account for difference in transfection efficiency. Cells were treated with NITD008, a DENV RNA-dependent RNA polymerase inhibitor, served as control. Statistical significance compared to Huh7 NT cells is indicated (n.s., non-significant). (**E**) The m^5^C1218 methylation level of DENV gRNA extracted from Huh7 NT and NSUN6 KO cells was measured by targeted Miseq bisulfite sequencing. Shown are methylation heatmaps of the cytosines in the amplicon nt 1106-1266 (n=1). (**F**) NSUN6 *in vitro* methylation assay (n=1). Recombinant NSUN6 was incubated with ^3^H-labeled SAM and 750 nM of different DENV IVT substrates (DENV gRNA IVT and DENV E-coding sequence IVT). An IVT of the eGFP-coding sequence was used as negative control. tRNA^Thr^ as was as positive control. Shown are the methylation levels (mean ± SD). Statistical significance is indicated on the top (p < 0.01; ***, p < 0.001; ****, p < 0.0001).

The m^5^C modification affects several biological processes (55). Therefore, we initially sought to assess the effect of NSUN6 depletion on processes that may indirectly influence the viral life cycle. First, we measured the levels of m^5^C on tRNA^Thr^, which is modified at position C72 by NSUN6 (54). Of note, tRNA^Thr^ is additionally methylated by NSUN2 in the tRNA variable loop (C48-50) (56,57). We used DORQ-seq, a hybridization-based approach that allows the purification of specific tRNAs (39), followed by LC-MS/MS (Fig. **3B**). The m^5^C levels on tRNA^Thr^ isolated from Huh7 NSUN6 KO cells were reduced by approximately 30% compared to that isolated from Huh7 NT cells. This result supported the role of NSUN6 in catalyzing one of three m^5^C sites on tRNA^Thr^ and suggested that NSUN6 deficiency did not alter the methylation of the other two sites by NSUN2. Moreover, depletion of NSUN6 had no significant effect on global translation (Fig. **3C**) or cell growth (Supplementary Fig. **S3C**). Next, we examined a potential effect of NSUN6 depletion on DENV gRNA replication. For this, Huh7 NT and NSUN6 KO cells were electroporated with a full-length DENV *firefly* luciferase reporter RNA to bypass the natural entry pathway of the virus. Luciferase activity was used as readout for viral replication levels over time. Cells treated with NITD008, an adenosine nucleoside inhibitor that potently abrogates DENV replication (41), served as negative control (Fig. **3D**). Similar replication levels were observed in Huh7 NT and NSUN6 KO cells, indicating that NSUN6 is dispensable for DENV gRNA replication (Fig. **3D**).

To determine the direct involvement of NSUN6 in DENV gRNA m^5^C1218 methylation, targeted MiSeq bisulfite sequencing was performed on total RNA extracted from infected Huh7 NT and NSUN6 KO cells (Fig. **3E**), as well as RNA extracted from virions released in the respective supernatants (Supplementary Fig. **S3D**). In line with our earlier findings in infected parental Huh7 cells (Fig. **2A** and **B**), m^5^C1218 was identified in the gRNA extracted from Huh7 NT cells and undetectable in the gRNA extracted from the corresponding virions. In contrast, this modification was undetectable in the gRNA extracted from Huh7 NSUN6 KO cells, indicating a pivotal role of NSUN6 in catalyzing m^5^C1218 in DENV gRNA in the cytoplasm.

Finally, we established an *in vitro* methylation assay using recombinant full-length NSUN6 and assessed the activity of NSUN6 on DENV RNA substrates (Fig. **3F**). Recombinant full-length NSUN6 (Supplementary Fig. **S3E**) and ^3^H-labeled S-adenosyl methionine (SAM) were incubated with 750 nM IVT of full-length DENV gRNA or IVT of DENV E-coding sequence. Equal molarity of tRNA^Thr^ was used as positive control. An IVT of the eGFP-coding sequence that did not contain the CUCCA consensus motif was used as a negative control. NSUN6 was able to methylate both DENV RNA substrates *in vitro* (Fig. **3F**). Interestingly, the shorter E IVT exhibited a 2-fold higher methylation level than the full-length gRNA IVT, suggesting that within the context of the full-length gRNA the C1218 site may be less accessible to the enzyme due to long-range RNA interactions, which may lead to tertiary structures (48,49).

### NSUN6-mediated m^5^C1218 destabilizes DENV gRNA

Among the different cellular processes, the m^5^C modification influences mRNA turnover by promoting either degradation or stabilization, depending on the target mRNA (58–60). We hypothesized a similar scenario for DENV gRNA and thus assessed the impact of NSUN6 on DENV gRNA turnover. Typically, the half-life (t_1/2_) of cellular RNAs determined by measuring RNA abundance after inhibition of transcription by actinomycin D treatment in time-course experiments. In contrast to cellular DNA-dependent RNA polymerases, DENV RNA-dependent RNA polymerase NS5 is not inhibited by actinomycin D. Thus, we adapted the approach and measured gRNA abundance following chemical inhibition of DENV gRNA synthesis using the NS5 inhibitor NITD008 (41) (see also Fig. **3D**). To ensure sufficient replication and detectable gRNA levels by qRT-PCR, Huh7 NT or NSUN6 KO cells were infected with DENV for 24 h before inhibition of NS5 with NITD008 treatment. Cells treated with DMSO served as control. Cells were then harvested at 24-hour intervals over a five-day period. DENV gRNA t_1/2_ increased from 40 h in Huh7 NT cells to approx. 100 h in Huh7 NSUN6 KO cells, indicating that NSUN6destabilizes DENV gRNA (Fig. **4A**). This effect was also reflected in the abundance of gRNA measured over time in the absence of NITD008 treatment. Huh7 NSUN6 KO cells accumulated more gRNA during infection than Huh7 NT cells (Fig. **4B**).

**Fig. 4.**
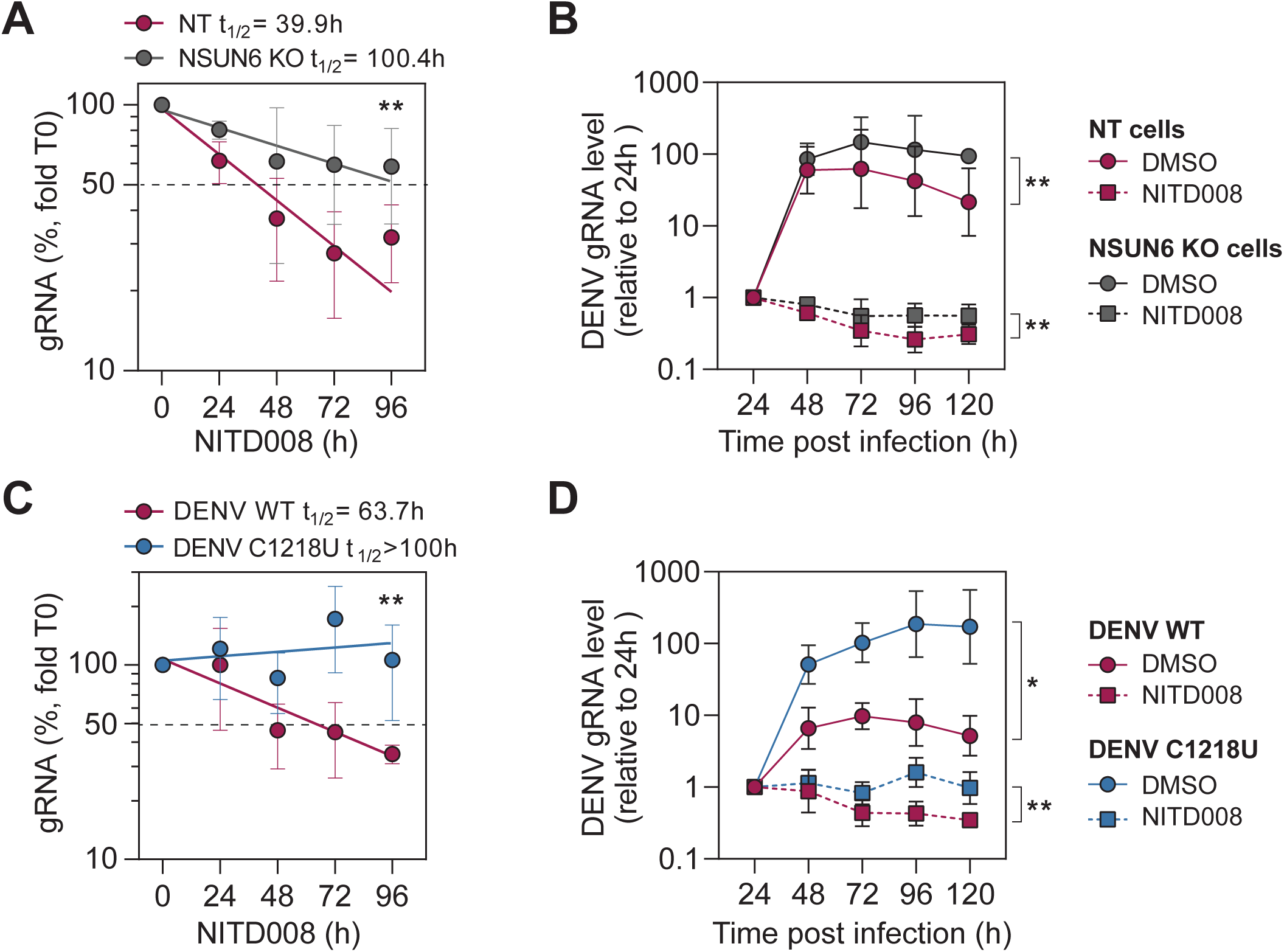
NSUN6-mediated m^5^C1218 destabilizes DENV gRNA. Measurement of DENV gRNA mRNA decay. (**A**) Huh7 NT and NSUN6 KO cells were infected with DENV for 24h before treatment with the DENV replication inhibitor NITD008. Cells were harvested at 24 h-interval and DENV gRNA abundance analyzed by Taqman probe-based qRT-PCR (n=3). Levels of DENV gRNA were normalized to *GAPDH* mRNA levels and to T_0_ (time of NITD008 addition). Shown are mean ± SD. Half-life (t_1/2_) of DENV gRNA was calculated. Statistical significance over all time points compared to Huh7 NT cells is indicated. **, p < 0.01. (**B**) DENV gRNA levels in Huh7 NT and NSUN6 KO cells infected for 120 h (n=3). Cells treated with NITD008 served as control. Cells were harvested at 24-h interval and DENV gRNA quantified by qRT-PCR (mean ± SD). Values were normalized to *GAPDH* mRNA levels and to time point 24 h post infection. Statistical significance over all time points compared to Huh7 NT cells is indicated. **, p < 0.01. (**C**) Decay measurement of DENV WT gRNA WT and DENV C1218U gRNA in Huh7 cells (mean ± SD, n=3). Statistical significance over all time points compared to DENV WT is indicated. **, p < 0.01. (**D**) DENV WT and DENV C1218U gRNA levels in Huh7 cells (mean ± SD, n=3). Statistical significance over all time points compared to DENV WT is indicated (*, p < 0.05; **, p < 0.01).

To establish a direct causal relationship between m^5^C1218 and DENV gRNA turnover, we generated a mutant virus containing a synonymous substitution at position 1218 (DENV C1218U) that disrupted the CUCCA motif. The half-lives of DENV wild type (WT) and DENV C1218U gRNAs were measured after infection of parental Huh7 cells, as previously established for the KO cells. Remarkably, the C1218U mutant gRNA was entirely stable over the 96 h time course, compared to a half-life of 63 h for the WT gRNA (Fig. **4C**). This result indicates that m^5^C1218 exerts a strong destabilizing effect on DENV gRNA. Stabilization of DENV C1218U gRNA also resulted in a pronounced accumulation of gRNA during infection (Fig. **4D**).

Taken together, our results demonstrate that NSUN6 catalyzes a highly specific m^5^C modification at position 1218 of DENV gRNA, which promotes DENV gRNA turnover and thereby strongly attenuates gRNA accumulation in DENV infected cells.

## DISCUSSION

In this study, we have established ViREn, an approach to purify and enrich viral RNAs, which is ideally suited for profiling of RNA modifications using RNA sequencing-based approaches. We used DENV, a (+)ss RNA virus as a model system, whose gRNA is over 10.7 kb long and not polyadenylated at its 3’ end. ViREn is a method that consists of two steps: 1) the removal of the most abundant and highly modified cellular RNAs based on the target RNA sedimentation coefficient in sucrose gradient ultracentrifugation, and 2) sequence-specific affinity capture of the target RNA. Despite modest yields, the results demonstrated a significant reduction in background signals for Illumina-based RNA-Seq and provided sufficient target RNA coverage to accurately identify RNA modifications in native gRNA molecules using DRS. Scale-up of production will allow the method to be combined with the use of LC-MS/MS to facilitate the detection of other RNA modifications not readily distinguishable with DRS using current basecaller algorithms. Finally, ViREn offers design versatility and straightforward transposability to other positive and negative-sense ssRNA viruses and should also be applicable to cellular target RNAs. In the future, it will facilitate the identification or validation of other RNA modifications such as m^6^A, inosine, Nm and pseudouridines.

Direct RNA sequencing of a 10.7 kb gRNA molecule with current nanopore technology remains challenging, especially for RNAs purified from cells. The ViREn approach proved essential for the reduction of cellular RNA and sfRNA contaminants, which would have otherwise occupied most of the sequencing pores and significantly limited the sequencing coverage of the DENV gRNA. Nevertheless, the success rate of passing the entire gRNA molecule through the sequencing pore was poor. Weobserved that the 3’ end stem-loop structure considerably hindered the entrance of the gRNA into the pore and a custom adapter directly targeting the 3’ end was not efficiently ligated. This issue was addressed on the one hand by the addition of a poly(A) tail and on the other hand by using complementary oligonucleotides to disrupt the secondary structures present in DENV 3’ UTR. The use of complementary oligonucleotides binding along the entire length of the gRNA led to improved sequencing throughput, presumably by disrupting more complex secondary and tertiary structures that appeared to be resistant to or able to refold after the various denaturation steps of the protocol.

Using stringent BSSeq, targeted MiSeq bisulfite sequencing and DRS tailored to DENV gRNA, we identified and validated the presence of a single m^5^C site at position 1218, in the E-coding sequence, in approximately 10-20% of DENV gRNA. Consistent with its life cycle being exclusively cytoplasmic, our findings demonstrated that m^5^C1218 is catalyzed by the host cytosolic methyltransferase NSUN6. As previously described for NSUN6 target mRNAs, the m^5^C modification occurred on the first cytosine of a CUCCA consensus sequence motif within the DENV gRNA sequence. Using Huh7 NSUN6 KO cells and targeted MiSeq bisulfite sequencing, we confirmed the direct involvement of NSUN6 in the modification of DENV gRNA m^5^C1218, with a 10-20% occurrence, similar to the median level of m^5^C methylation in human mRNAs (50,51). It is noteworthy that although *in silico* analysis of the DENV gRNA sequence revealed the presence of 16 CUCCA consensus sequence motifs, including one in the 3’ UTR (52), only one was modified by NSUN6 in our experiments. Furthermore, while the position of this motif in the E-coding sequence is conserved *in silico* in other DENV serotype 2 strains, e.g. 16681, it is not conserved in the other three DENV serotypes. The DENV gRNA architecture is complex. SHAPE analyses of DENV gRNAs purified from viral particles have revealed numerous secondary and tertiary structures that vary between serotypes, as well as long-range interactions that are important for viral replication and virus fitness (48,49,61,62). We assume that the CUCCA motif at C1218 in DENV2 NGC gRNA may be more accessible to the methyltransferase due to lack of secondary structure, although other factors, such as nearby binding sites for RNA-binding proteins, may also contribute to modification selectivity. Whether m^5^C modifications are present in other regions of the gRNA in different DENV serotypes and other flaviviruses will have to be investigated in the future.

The m^5^C modification plays a role in several post-transcriptional RNA processes, including nuclear export, splicing, stability, and translation of mRNAs (55). The deposition of m^5^C is dynamic, and effects likely depend on the reader proteins that will recognize the modification (55). These include Aly/REF export factor (ALYREF) (50), Y-Box binding protein 1 (YBX1) (58), LIN28B (63), YTH domain-containing family 2 (YTHDF2) (64) and serine/arginine-rich splicing factor 2 (SRSF2) (65). For example, ALYREF promotes the nuclear export of the regulatory-associated protein of mTOR *RPTOR* mRNA (66), and YBX1 stabilizes the heparin binding growth factor *HBGF* mRNA (58). Conversely, the m^5^C modification can also enhance mRNA decay, as observed for interferon regulatory factor 3 *IRF3* mRNA, resulting in a dampened interferon response during various viral infections (67). By using a mutant virus with a synonymous substitution at position C1218, we have shown that this single NSUN6-mediate m^5^C modification promotes the degradation of the gRNA by a yet unknown mechanism. It is interesting to note that YBX1 was reported to act at different steps of the DENV viral cycle with opposite effects. While its binding to the 3’ UTR negatively regulates viral replication (68), the interactions of YBX1 with the capsid and E proteins are necessary for the late steps of the viral life cycle, during the assembly of the viral particles (69,70). The identification of the m^5^C reader protein will be important to elucidate the mechanism of DENV gRNA turnover.

Emerging evidence indicates that NSUN2 methylates a variety of viral RNAs. Hepatitis B virus, a DNA virus, replicates in the nucleus where its viral mRNAs are modified by NSUN2 at position 1291. The modification is recognized by ALYREF, which promotes export and translation of the viral mRNAs (71). Recently, NSUN2 has also been reported to partially localize to the cytosol during infection with several (+)ssRNA viruses, where it modifies viral gRNAs. In HCV, a virus closely related to DENV, a single m^5^C modification at position 7525 is recognized by YBX1 and increases gRNA stability and viral replication (72,73). In Enterovirus 71, multiple m^5^C modifications increase gRNA stability and translation (74). Conversely, the various m^5^C modifications identified in the gRNA of severe acute respiratory syndrome coronavirus 2 promote its decay (75). However, these results are being challenged (14) as the methods and rigor of the analyses improve.

Aligning with this observation and previously described limitations of RNA modification-specific antibody immunoprecipitation approaches (76,77), our analysis of DENV gRNA DRS with Dorado 0.8.2, which also provides models for m^6^A, did not identify any m^6^A sites (Supplementary Table **S2**), also agreeing with recent antibody-independent SELECT analyses (13). Taken together, these discrepancies underscore the importance of orthogonal validation to ascertain the presence of RNA modifications identified as well as their functional validation.

While the m^5^C-modified gRNAs of the viruses mentioned above were also found in viral particles, the NSUN6-mediated m^5^C1218 modification in DENV gRNA was exclusively detected in cytoplasmic gRNAs, but not in virions. This observation raises the question of a potential sorting mechanism that could prevent gRNAs from being packaged into viral particles and target them for degradation. Currently, we have only a limited understanding of the mechanisms that regulate the fate of the gRNAs during the viral life cycle. One might be tempted to speculate that RNA modifications could mark a gRNA for replication, translation, degradation, or packaging. How DENV gRNA m^5^C1218 is recognized, and the role of this modification in the DENV life cycle, remains to be elucidated.

## ACKNOWLEDGEMENTS

We thank Sabine Wegehingel and Walter Nickel (Center for Biochemistry BZH, University Heidelberg, Germany) for providing help and access to the IncuCyte. We also thank Laurent Chatel-Chaix (Centre Armand-Frappier Santé Biotechnologie, Canada) for critical reading of the manuscript. We thank the Institute of Genetics and Biophysics A. Buzzati-Traverso, The National Research Council of Italy for the support to Francesca Tuorto.

## AUTHOR CONTRIBUTIONS

A.R. and M.H conceptualized the study; C.C.W., I.S.N., J.H., Z.N., K.K., V.M. Z.Ö, D.L., S.S. and S.F. performed and analyzed experiments under the supervision of M.F., T.S., G.S., Y.M., F.T., C.D., M.H. and A.R.; J.S. analyzed the RNA-Seq data. M.F., N.P., T.S., G.S., Y.M., F.T., C.D., M.H. and A.R.; C.C.W. and A.R. made the original draft of the manuscript. All authors edited and approved the final manuscript.

## SUPPLEMENTARY FIGURE LEGENDS

**Supplementary Fig. S1.**
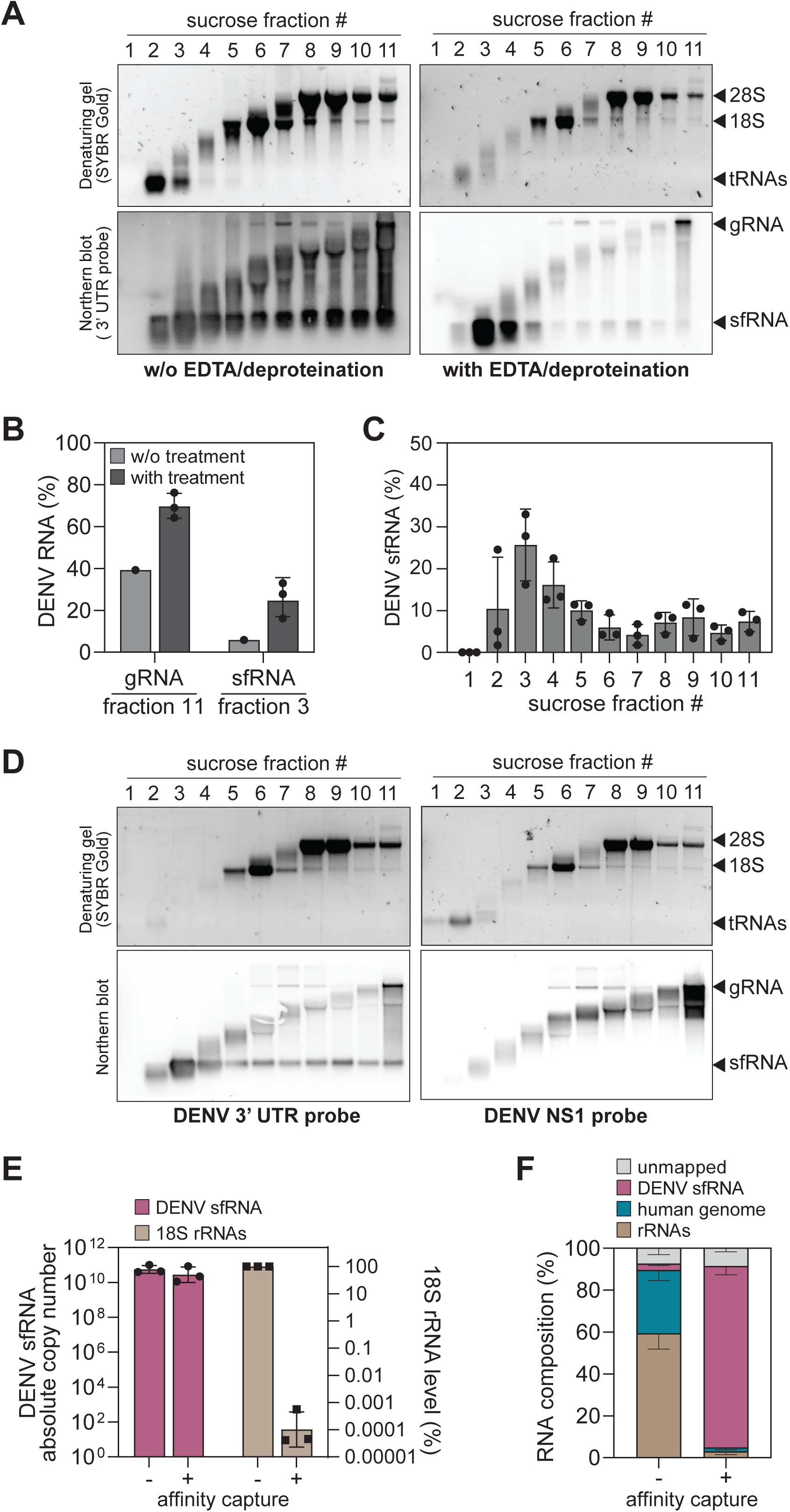
Establishment of the ViREn method. (**A** and **B**) Total RNA extracted from infected cells separated by sucrose gradient ultracentrifugation. The distribution of cellular RNAs and DENV RNAs extracted from infected cells across sucrose density gradient fractions was detected using a denaturing agarose gel and corresponding northern blot using a hybridization probe targeting DENV 3’ UTR. (A) Before separation, the total RNA was isolated using acid-guanidinium-phenol-chloroform extraction (left panel) or additionally treated with EDTA and deproteinated (right panel). (B) Quantification of the corresponding gRNA distribution in fraction 11 and sfRNA in fraction 3. (**C**) Quantification of DENV sfRNA detected in each sucrose gradient fraction from infected cells (mean ± SD, n=3). (**D**) The distribution of cellular RNAs and DENV RNAs extracted from infected cells across sucrose density gradient fractions was detected using a denaturing agarose gel. Corresponding northern blots were labeled using a hybridization probe targeting DENV 3’ UTR (left panel) or NS1-coding region (right panel). (**E**) DENV sfRNA collected in fraction 3 was further purified by affinity capture. The level of DENV sfRNA and 18S rRNA contaminant, before and after affinity capture, was measured by Taqman probe-based qRT-PCR (mean ± SD, n=3). (**F**) The total RNA content of fraction 3 before and after affinity capture assay was analyzed by Illumina-based RNA-Seq (n=3). Shown is the proportion of reads in each sample corresponding to DENV sfRNA, human genome, rRNAs, and unmapped reads (mean ± SD, n=3).

**Supplementary Fig. S2.**
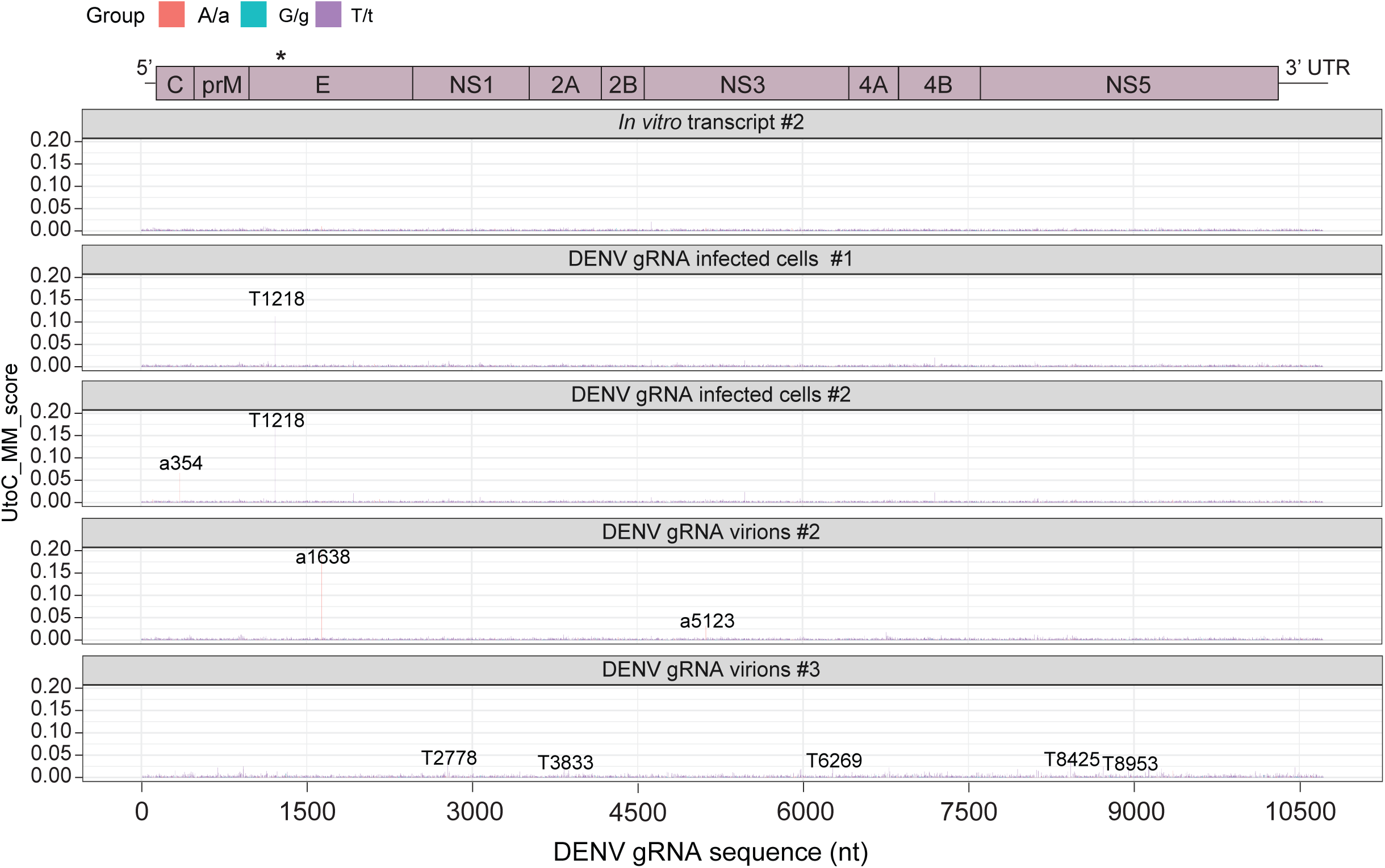
Mapping of m^5^C on DENV gRNA BSSeq analysis. DENV gRNA purified from infected cells and virions were subjected to BSSeq (n=3). Unmethylated *in vitro* transcript processed with the ViREn method was used for comparative analysis and employed to reduce background noise. Shown are two biological replicates. The third is shown in Fig. 2. The asterisk marks the m^5^C position at nt 1218 in the E-coding sequence of DENV gRNA. A schematic of DENV gRNA organization is shown on the top.

**Supplementary Fig. S3.**
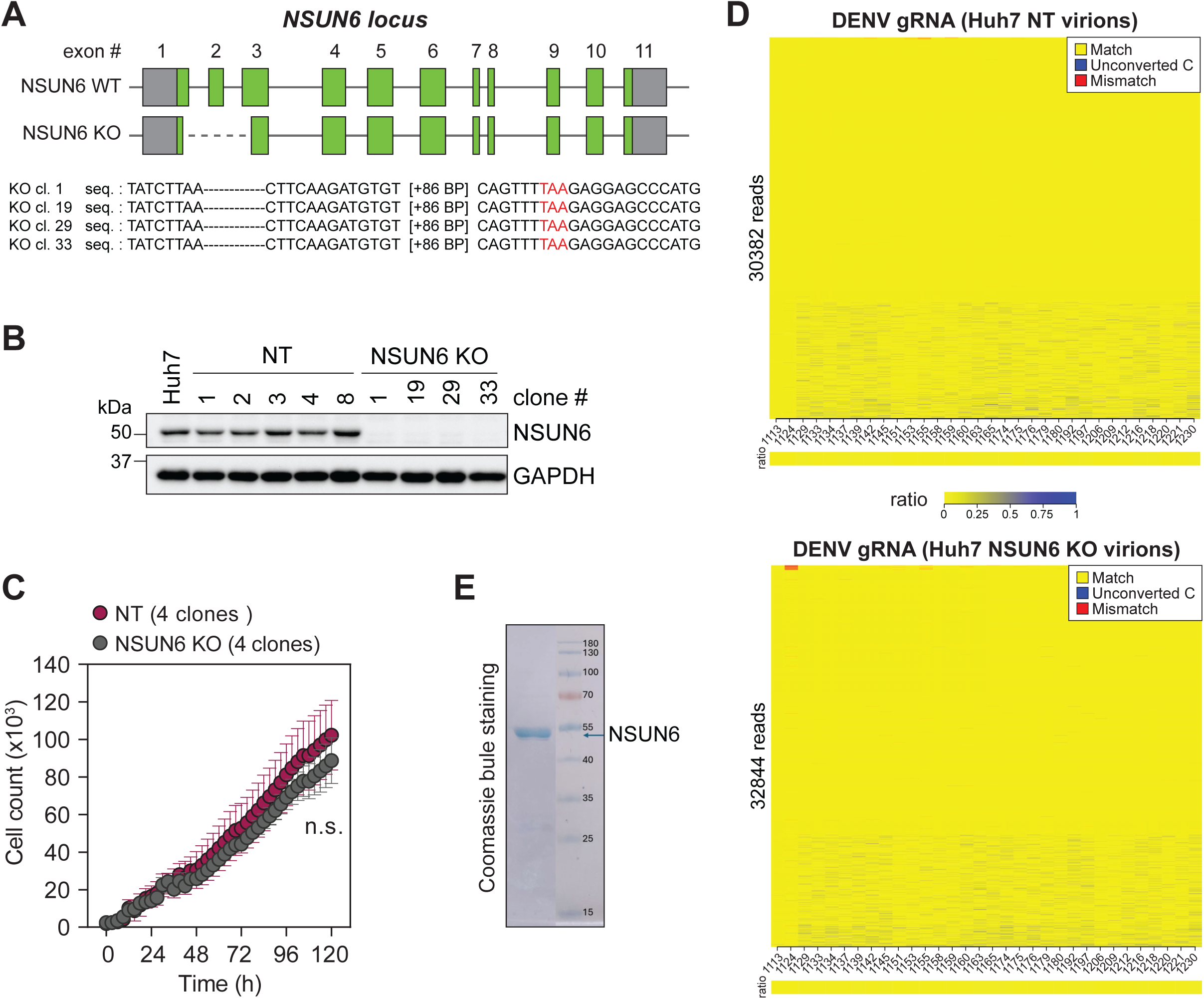
Characterization of Huh7 NSUN6 KO cell clones. (**A**) Schematics of *NSUN6* locus edited using the two-guide CRISPR-Cas9 approach. NSUN6 open reading frame (ORF) is indicated in green. Huh7 NSUN6 KO cells were generated by creating a deletion in the N-terminus of the ORF using two guide RNAs targeting *NSUN6* exon 1 and 3. Four homozygous clones were selected based on the deletion of the genomic DNA sequence between the guide RNAs and confirmed by sequencing. Sequences for each clone are indicated. Nucleotide deletions are indicated by black dashes. The appearance of a premature stop codon is indicated in red. (**B**) Representative immunoblot of NSUN6 protein expression levels in parental Huh7 cells, non-targeting (NT) cell clones, and NSUN6 KO cell clones. GADPH served as loading control. (**C**) Cells growth of Huh7 NT and NSUN6 cell clones was monitored at 3-h interval for 5 days using the IncuCyte live-cell analysis system. Shown is the number of cells. Statistical significance over all time points compared to Huh7 NT cell clones is indicated (n.s., non-significant). (**D**) The m^5^C1218 methylation level of DENV gRNA extracted from Huh7 NT and NSUN6 KO virions was measured by targeted Miseq bisulfite sequencing. Shown are methylation heatmaps of the cytosines in the amplicon nt 1106-1266 (n=1). (**E**) Purification of recombinant full-length NSUN6. Shown is a Coomassie staining.

## Footnotes

1 https://www.ecdc.europa.eu/en/publications-data/twelve-month-dengue-virus-disease-case-notification-rate-100-000-population

## REFERENCES

1. Helm, M. and Motorin, Y. (2017) Detecting RNA modifications in the epitranscriptome: predict and validate. Nat Rev Genet, 18, 275–291.

2. Cappannini, A., Ray, A., Purta, E., Mukherjee, S., Boccaletto, P., Moafinejad, S.N., Lechner, A., Barchet, C., Klaholz, B.P., Stefaniak, F. et al. (2024) MODOMICS: a database of RNA modifications and related information. 2023 update. Nucleic Acids Res, 52, D239–D244.

3. McCown, P.J., Ruszkowska, A., Kunkler, C.N., Breger, K., Hulewicz, J.P., Wang, M.C., Springer, N.A. and Brown, J.A. (2020) Naturally occurring modified ribonucleosides. Wiley Interdiscip Rev RNA, 11, e1595.

4. Sun, H., Li, K., Liu, C. and Yi, C. (2023) Regulation and functions of non-m(6)A mRNA modifications. Nat Rev Mol Cell Biol, 24, 714–731.

5. Boo, S.H. and Kim, Y.K. (2020) The emerging role of RNA modifications in the regulation of mRNA stability. Exp Mol Med, 52, 400–408.

6. Huang, G., Zhang, F., Xie, D., Ma, Y., Wang, P., Cao, G., Chen, L., Lin, S., Zhao, Z. and Cai, Z. (2023) High-throughput profiling of RNA modifications by ultra-performance liquid chromatography coupled to complementary mass spectrometry: Methods, quality control, and applications. Talanta, 263, 124697.

7. Herbert, C., Valesyan, S., Kist, J. and Limbach, P.A. (2024) Analysis of RNA and Its Modifications. Annu Rev Anal Chem (Palo Alto Calif), 17, 47–68.

8. Garalde, D.R., Snell, E.A., Jachimowicz, D., Sipos, B., Lloyd, J.H., Bruce, M., Pantic, N., Admassu, T., James, P., Warland, A. et al. (2018) Highly parallel direct RNA sequencing on an array of nanopores. Nat Methods, 15, 201–206.

9. Workman, R.E., Tang, A.D., Tang, P.S., Jain, M., Tyson, J.R., Razaghi, R., Zuzarte, P.C., Gilpatrick, T., Payne, A., Quick, J. et al. (2019) Nanopore native RNA sequencing of a human poly(A) transcriptome. Nat Methods, 16, 1297–1305.

10. Gokhale, N.S., McIntyre, A.B.R., McFadden, M.J., Roder, A.E., Kennedy, E.M., Gandara, J.A., Hopcraft, S.E., Quicke, K.M., Vazquez, C., Willer, J. et al. (2016) N6-Methyladenosine in Flaviviridae Viral RNA Genomes Regulates Infection. Cell Host Microbe, 20, 654–665.

11. Lichinchi, G., Zhao, B.S., Wu, Y., Lu, Z., Qin, Y., He, C. and Rana, T.M. (2016) Dynamics of Human and Viral RNA Methylation during Zika Virus Infection. Cell Host Microbe, 20, 666–673.

12. McIntyre, W., Netzband, R., Bonenfant, G., Biegel, J.M., Miller, C., Fuchs, G., Henderson, E., Arra, M., Canki, M., Fabris, D. et al. (2018) Positive-sense RNA viruses reveal the complexity and dynamics of the cellular and viral epitranscriptomes during infection. Nucleic Acids Res, 46, 5776–5791.

13. Baquero-Perez, B., Yonchev, I.D., Delgado-Tejedor, A., Medina, R., Puig-Torrents, M., Sudbery, I., Begik, O., Wilson, S.A., Novoa, E.M. and Diez, J. (2024) N(6)-methyladenosine modification is not a general trait of viral RNA genomes. Nat Commun, 15, 1964.

14. Huang, A., Riepler, L., Rieder, D., Kimpel, J. and Lusser, A. (2023) No evidence for epitranscriptomic m(5)C modification of SARS-CoV-2, HIV and MLV viral RNA. RNA, 29, 756–763.

15. Mazeaud, C., Freppel, W. and Chatel-Chaix, L. (2018) The Multiples Fates of the Flavivirus RNA Genome During Pathogenesis. Front Genet, 9, 595.

16. Jones, C.I., Zabolotskaya, M.V. and Newbury, S.F. (2012) The 5’ --> 3’ exoribonuclease XRN1/Pacman and its functions in cellular processes and development. Wiley Interdiscip Rev RNA, 3, 455–468.

17. Chapman, E.G., Costantino, D.A., Rabe, J.L., Moon, S.L., Wilusz, J., Nix, J.C. and Kieft, J.S. (2014) The structural basis of pathogenic subgenomic flavivirus RNA (sfRNA) production. Science, 344, 307–310.

18. Chapman, E.G., Moon, S.L., Wilusz, J. and Kieft, J.S. (2014) RNA structures that resist degradation by Xrn1 produce a pathogenic Dengue virus RNA. Elife, 3, e01892.

19. Cumberworth, S.L., Clark, J.J., Kohl, A. and Donald, C.L. (2017) Inhibition of type I interferon induction and signalling by mosquito-borne flaviviruses. Cell Microbiol, 19.

20. Manokaran, G., Finol, E., Wang, C., Gunaratne, J., Bahl, J., Ong, E.Z., Tan, H.C., Sessions, O.M., Ward, A.M., Gubler, D.J. et al. (2015) Dengue subgenomic RNA binds TRIM25 to inhibit interferon expression for epidemiological fitness. Science, 350, 217–221.

21. Moon, S.L., Dodd, B.J., Brackney, D.E., Wilusz, C.J., Ebel, G.D. and Wilusz, J. (2015) Flavivirus sfRNA suppresses antiviral RNA interference in cultured cells and mosquitoes and directly interacts with the RNAi machinery. Virology, 485, 322–329.

22. Slonchak, A. and Khromykh, A.A. (2018) Subgenomic flaviviral RNAs: What do we know after the first decade of research. Antiviral Res, 159, 13–25.

23. Zhang, Z., Jiang, L. and Zeng, G. (2018) Non-coding RNA: a key regulator of the pathogenicity and immunity of Flaviviridae viruses infection. Cell Mol Immunol, 15, 185–186.

24. Pijlman, G.P., Funk, A., Kondratieva, N., Leung, J., Torres, S., van der Aa, L., Liu, W.J., Palmenberg, A.C., Shi, P.Y., Hall, R.A., et al. (2008) A highly structured, nuclease-resistant, noncoding RNA produced by flaviviruses is required for pathogenicity. Cell Host Microbe, 4, 579–591.

25. Liu-Wei, W., van der Toorn, W., Bohn, P., Holzer, M., Smyth, R.P. and von Kleist, M. (2024) Sequencing accuracy and systematic errors of nanopore direct RNA sequencing. BMC Genomics, 25, 528.

26. Gualano, R.C., Pryor, M.J., Cauchi, M.R., Wright, P.J. and Davidson, A.D. (1998) Identification of a major determinant of mouse neurovirulence of dengue virus type 2 using stably cloned genomic-length cDNA. J Gen Virol, 79 (Pt 3), 437–446.

27. Kumar, A., Buhler, S., Selisko, B., Davidson, A., Mulder, K., Canard, B., Miller, S. and Bartenschlager, R. (2013) Nuclear localization of dengue virus nonstructural protein 5 does not strictly correlate with efficient viral RNA replication and inhibition of type I interferon signaling. J Virol, 87, 4545–4557.

28. Roth, H., Magg, V., Uch, F., Mutz, P., Klein, P., Haneke, K., Lohmann, V., Bartenschlager, R., Fackler, O.T., Locker, N. et al. (2017) Flavivirus Infection Uncouples Translation Suppression from Cellular Stress Responses. mBio, 8.

29. van den Hoff, M.J., Christoffels, V.M., Labruyere, W.T., Moorman, A.F. and Lamers, W.H. (1995) Electrotransfection with “intracellular” buffer. Methods Mol Biol, 48, 185–197.

30. Henchal, E.A., Gentry, M.K., McCown, J.M. and Brandt, W.E. (1982) Dengue virus-specific and flavivirus group determinants identified with monoclonal antibodies by indirect immunofluorescence. Am J Trop Med Hyg, 31, 830–836.

31. Langmead, B., Trapnell, C., Pop, M. and Salzberg, S.L. (2009) Ultrafast and memory-efficient alignment of short DNA sequences to the human genome. Genome Biol, 10, R25.

32. Dobin, A., Davis, C.A., Schlesinger, F., Drenkow, J., Zaleski, C., Jha, S., Batut, P., Chaisson, M. and Gingeras, T.R. (2013) STAR: ultrafast universal RNA-seq aligner. Bioinformatics, 29, 15–21.

33. Liao, Y., Smyth, G.K. and Shi, W. (2014) featureCounts: an efficient general purpose program for assigning sequence reads to genomic features. Bioinformatics, 30, 923–930.

34. Thorvaldsdottir, H., Robinson, J.T. and Mesirov, J.P. (2013) Integrative Genomics Viewer (IGV): high-performance genomics data visualization and exploration. Brief Bioinform, 14, 178–192.

35. Dai, Q., Ye, C., Irkliyenko, I., Wang, Y., Sun, H.L., Gao, Y., Liu, Y., Beadell, A., Perea, J., Goel, A. et al. (2024) Ultrafast bisulfite sequencing detection of 5-methylcytosine in DNA and RNA. Nat Biotechnol, 42, 1559–1570.

36. Bormann, F., Tuorto, F., Cirzi, C., Lyko, F. and Legrand, C. (2019) BisAMP: A web-based pipeline for targeted RNA cytosine-5 methylation analysis. Methods, 156, 121–127.

37. Baek, A., Lee, G.E., Golconda, S., Rayhan, A., Manganaris, A.A., Chen, S., Tirumuru, N., Yu, H., Kim, S., Kimmel, C. et al. (2024) Single-molecule epitranscriptomic analysis of full-length HIV-1 RNAs reveals functional roles of site-specific m(6)As. Nat Microbiol, 9, 1340–1355.

38. Magg, V., Manetto, A., Kopp, K., Wu, C.C., Naghizadeh, M., Lindner, D., Eke, L., Welsch, J., Kallenberger, S.M., Schott, J. et al. (2024) Turnover of PPP1R15A mRNA encoding GADD34 controls responsiveness and adaptation to cellular stress. Cell Rep, 43, 114069.

39. Kristen, M., Lander, M., Kilz, L.M., Gleue, L., Jorg, M., Bregeon, D., Hamdane, D., Marchand, V., Motorin, Y., Friedland, K. et al. (2024) DORQ-seq: high-throughput quantification of femtomol tRNA pools by combination of cDNA hybridization and Deep sequencing. Nucleic Acids Res, 52, e89.

40. Kellner, S., Ochel, A., Thuring, K., Spenkuch, F., Neumann, J., Sharma, S., Entian, K.D., Schneider, D. and Helm, M. (2014) Absolute and relative quantification of RNA modifications via biosynthetic isotopomers. Nucleic Acids Res, 42, e142.

41. Yin, Z., Chen, Y.L., Schul, W., Wang, Q.Y., Gu, F., Duraiswamy, J., Kondreddi, R.R., Niyomrattanakit, P., Lakshminarayana, S.B., Goh, A. et al. (2009) An adenosine nucleoside inhibitor of dengue virus. Proc Natl Acad Sci U S A, 106, 20435–20439.

42. Livak, K.J. and Schmittgen, T.D. (2001) Analysis of relative gene expression data using real-time quantitative PCR and the 2(-Delta Delta C(T)) Method. Methods, 25, 402–408.

43. Stollar, V., Stevens, T.M. and Schlesinger, R.W. (1966) Studies on the nature of dengue viruses. II. Characterization of viral RNA and effects of inhibitors of RNA synthesis. Virology, 30, 303–312.

44. Cleaves, G.R., Ryan, T.E. and Schlesinger, R.W. (1981) Identification and characterization of type 2 dengue virus replicative intermediate and replicative form RNAs. Virology, 111, 73–83.

45. Lu, L., Zhang, X., Zhou, Y., Shi, Z., Xie, X., Zhang, X., Gao, L., Fu, A., Liu, C., He, B. et al. (2024) Base-resolution m(5)C profiling across the mammalian transcriptome by bisulfite-free enzyme-assisted chemical labeling approach. Mol Cell, 84, 2984–3000 e2988.

46. Schaefer, M., Pollex, T., Hanna, K. and Lyko, F. (2009) RNA cytosine methylation analysis by bisulfite sequencing. Nucleic Acids Res, 37, e12.

47. Motorin, Y., Lyko, F. and Helm, M. (2010) 5-methylcytosine in RNA: detection, enzymatic formation and biological functions. Nucleic Acids Res, 38, 1415–1430.

48. Dethoff, E.A., Boerneke, M.A., Gokhale, N.S., Muhire, B.M., Martin, D.P., Sacco, M.T., McFadden, M.J., Weinstein, J.B., Messer, W.B., Horner, S.M. et al. (2018) Pervasive tertiary structure in the dengue virus RNA genome. Proc Natl Acad Sci U S A, 115, 11513–11518.

49. Boerneke, M.A., Gokhale, N.S., Horner, S.M. and Weeks, K.M. (2023) Structure-first identification of RNA elements that regulate dengue virus genome architecture and replication. Proc Natl Acad Sci U S A, 120, e2217053120.

50. Yang, X., Yang, Y., Sun, B.F., Chen, Y.S., Xu, J.W., Lai, W.Y., Li, A., Wang, X., Bhattarai, D.P., Xiao, W. et al. (2017) 5-methylcytosine promotes mRNA export - NSUN2 as the methyltransferase and ALYREF as an m(5)C reader. Cell Res, 27, 606–625.

51. Huang, T., Chen, W., Liu, J., Gu, N. and Zhang, R. (2019) Genome-wide identification of mRNA 5-methylcytosine in mammals. Nat Struct Mol Biol, 26, 380–388.

52. Selmi, T., Hussain, S., Dietmann, S., Heiss, M., Borland, K., Flad, S., Carter, J.M., Dennison, R., Huang, Y.L., Kellner, S. et al. (2021) Sequence- and structure-specific cytosine-5 mRNA methylation by NSUN6. Nucleic Acids Res, 49, 1006–1022.

53. Liu, J., Huang, T., Zhang, Y., Zhao, T., Zhao, X., Chen, W. and Zhang, R. (2021) Sequence- and structure-selective mRNA m(5)C methylation by NSUN6 in animals. Natl Sci Rev, 8, nwaa273.

54. Haag, S., Warda, A.S., Kretschmer, J., Gunnigmann, M.A., Hobartner, C. and Bohnsack, M.T. (2015) NSUN6 is a human RNA methyltransferase that catalyzes formation of m5C72 in specific tRNAs. RNA, 21, 1532–1543.

55. Chen, Y.S., Yang, W.L., Zhao, Y.L. and Yang, Y.G. (2021) Dynamic transcriptomic m(5) C and its regulatory role in RNA processing. Wiley Interdiscip Rev RNA, 12, e1639.

56. Van Haute, L., Lee, S.Y., McCann, B.J., Powell, C.A., Bansal, D., Vasiliauskaite, L., Garone, C., Shin, S., Kim, J.S., Frye, M., et al. (2019) NSUN2 introduces 5-methylcytosines in mammalian mitochondrial tRNAs. Nucleic Acids Res, 47, 8720–8733.

57. Khoddami, V. and Cairns, B.R. (2013) Identification of direct targets and modified bases of RNA cytosine methyltransferases. Nat Biotechnol, 31, 458–464.

58. Chen, X., Li, A., Sun, B.F., Yang, Y., Han, Y.N., Yuan, X., Chen, R.X., Wei, W.S., Liu, Y., Gao, C.C. et al. (2019) 5-methylcytosine promotes pathogenesis of bladder cancer through stabilizing mRNAs. Nat Cell Biol, 21, 978–990.

59. Wu, R., Sun, C., Chen, X., Yang, R., Luan, Y., Zhao, X., Yu, P., Luo, R., Hou, Y., Tian, R. et al. (2024) NSUN5/TET2-directed chromatin-associated RNA modification of 5-methylcytosine to 5-hydroxymethylcytosine governs glioma immune evasion. Proc Natl Acad Sci U S A, 121, e2321611121.

60. Guarnacci, M., Zhang, P.H., Kanchi, M., Hung, Y.T., Lin, H., Shirokikh, N.E., Yang, L. and Preiss, T. (2024) Substrate diversity of NSUN enzymes and links of 5-methylcytosine to mRNA translation and turnover. Life Sci Alliance, 7.

61. de Borba, L., Villordo, S.M., Iglesias, N.G., Filomatori, C.V., Gebhard, L.G. and Gamarnik, A.V. (2015) Overlapping local and long-range RNA-RNA interactions modulate dengue virus genome cyclization and replication. J Virol, 89, 3430–3437.

62. Huber, R.G., Lim, X.N., Ng, W.C., Sim, A.Y.L., Poh, H.X., Shen, Y., Lim, S.Y., Sundstrom, K.B., Sun, X., Aw, J.G. et al. (2019) Structure mapping of dengue and Zika viruses reveals functional long-range interactions. Nat Commun, 10, 1408.

63. Su, J., Wu, G., Ye, Y., Zhang, J., Zeng, L., Huang, X., Zheng, Y., Bai, R., Zhuang, L., Li, M. et al. (2021) NSUN2-mediated RNA 5-methylcytosine promotes esophageal squamous cell carcinoma progression via LIN28B-dependent GRB2 mRNA stabilization. Oncogene, 40, 5814–5828.

64. Dai, X., Gonzalez, G., Li, L., Li, J., You, C., Miao, W., Hu, J., Fu, L., Zhao, Y., Li, R. et al. (2020) YTHDF2 Binds to 5-Methylcytosine in RNA and Modulates the Maturation of Ribosomal RNA. Anal Chem, 92, 1346–1354.

65. Ma, H.L., Bizet, M., Soares Da Costa, C., Murisier, F., de Bony, E.J., Wang, M.K., Yoshimi, A., Lin, K.T., Riching, K.M., Wang, X., et al. (2023) SRSF2 plays an unexpected role as reader of m(5)C on mRNA, linking epitranscriptomics to cancer. Mol Cell, 83, 4239–4254 e4210.

66. Zhong, L., Wu, J., Zhou, B., Kang, J., Wang, X., Ye, F. and Lin, X. (2024) ALYREF recruits ELAVL1 to promote colorectal tumorigenesis via facilitating RNA m5C recognition and nuclear export. NPJ Precis Oncol, 8, 243.

67. Wang, H., Feng, J., Zeng, C., Liu, J., Fu, Z., Wang, D., Wang, Y., Zhang, L., Li, J., Jiang, A. et al. (2023) NSUN2-mediated M(5)c methylation of IRF3 mRNA negatively regulates type I interferon responses during various viral infections. Emerg Microbes Infect, 12, 2178238.

68. Paranjape, S.M. and Harris, E. (2007) Y box-binding protein-1 binds to the dengue virus 3’-untranslated region and mediates antiviral effects. J Biol Chem, 282, 30497–30508.

69. Diosa-Toro, M., Kennedy, D.R., Chuo, V., Popov, V.L., Pompon, J. and Garcia-Blanco, M.A. (2022) Y-Box Binding Protein 1 Interacts with Dengue Virus Nucleocapsid and Mediates Viral Assembly. mBio, 13, e0019622.

70. Phillips, S.L., Soderblom, E.J., Bradrick, S.S. and Garcia-Blanco, M.A. (2016) Identification of Proteins Bound to Dengue Viral RNA In Vivo Reveals New Host Proteins Important for Virus Replication. mBio, 7, e01865–01815.

71. Ding, S., Liu, H., Liu, L., Ma, L., Chen, Z., Zhu, M., Liu, L., Zhang, X., Hao, H., Zuo, L. et al. (2024) Epigenetic addition of m(5)C to HBV transcripts promotes viral replication and evasion of innate antiviral responses. Cell Death Dis, 15, 39.

72. Li, Z.L., Xie, Y., Wang, Y., Wang, J., Zhou, X. and Zhang, X.L. (2025) NSUN2-mediated HCV RNA m5C Methylation Facilitates Viral RNA Stability and Replication. Genomics Proteomics Bioinformatics.

73. Li, Z.L., Xie, Y., Xie, Y., Chen, H., Zhou, X., Liu, M. and Zhang, X.L. (2024) HCV 5-Methylcytosine Enhances Viral RNA Replication through Interaction with m5C Reader YBX1. ACS Chem Biol, 19, 1648–1660.

74. Liu, L., Chen, Z., Zhang, K., Hao, H., Ma, L., Liu, H., Yu, B., Ding, S., Zhang, X., Zhu, M. et al. (2024) NSUN2 mediates distinct pathways to regulate enterovirus 71 replication. Virol Sin, 39, 574–586.

75. Wang, H., Feng, J., Fu, Z., Xu, T., Liu, J., Yang, S., Li, Y., Deng, J., Zhang, Y., Guo, M. et al. (2024) Epitranscriptomic m(5)C methylation of SARS-CoV-2 RNA regulates viral replication and the virulence of progeny viruses in the new infection. Sci Adv, 10, eadn9519.

76. McIntyre, A.B.R., Gokhale, N.S., Cerchietti, L., Jaffrey, S.R., Horner, S.M. and Mason, C.E. (2020) Limits in the detection of m(6)A changes using MeRIP/m(6)A-seq. Sci Rep, 10, 6590.

77. Helm, M., Lyko, F. and Motorin, Y. (2019) Limited antibody specificity compromises epitranscriptomic analyses. Nat Commun, 10, 5669.

